# Aggresomes protect mRNA under stress in *Escherichia coli*

**DOI:** 10.1101/2024.04.27.591437

**Authors:** Linsen Pei, Yujia Xian, Xiaodan Yan, Charley Schaefer, Aisha H. Syeda, Jamieson A. L. Howard, Wei Zhang, Hebin Liao, Fan Bai, Mark C. Leake, Yingying Pu

**Affiliations:** The State Key Laboratory Breeding Base of Basic Science of Stomatology & Key Laboratory of Oral Biomedicine Ministry of Education, School & Hospital of Stomatology, Medical Research Institute, Wuhan University, China; Frontier Science Centre for Immunology and Metabolism, Wuhan University, China; School of Physics, Engineering and Technology, University of York, York, UK; Department of Biology, University of York, York, UK; Biomedical Pioneering Innovation Centre (BIOPIC), Peking-Tsinghua Center for Life Sciences (CLS), School of Life Sciences, Peking University, Beijing, China; State Key Laboratory of Metabolic Dysregulation & Prevention and Treatment of Esophageal Cancer, Biomedical Pioneering Innovation Center (BIOPIC), Peking University, Beijing, China; Peking University Beijing-Tianjin-Hebei Biomedical Pioneering Innovation Center, Tianjin, China; Department of Immunology, Hubei Province Key Laboratory of Allergy and Immunology, State Key Laboratory of Virology and Medical Research Institute, Wuhan University School of Basic Medical Sciences, China

## Abstract

Membraneless droplets formed through liquid-liquid phase separation of ribonucleoprotein particles contribute to mRNA storage in eukaryotic cells. How such aggresomes contribute to mRNA dynamics under stress, and their functional role, is less understood in bacteria. Here we used multiple approaches including live-cell imaging, polymer physics modelling and transcriptomics to show that prolonged stress leading to ATP depletion in *Escherichia coli* results in increased aggresome formation, compaction, and selective mRNA enrichment within these aggresomes. Longer transcripts accumulate more in aggresomes than in the cytosol.

Mass spectrometry and mutagenesis studies showed that mRNA ribonucleases are excluded from aggresomes due to electrostatic repulsion arising from their negative surface charges. Experiments with fluorescent reporters and disruption of aggresome formation showed that mRNA storage within aggresomes promoted rapid translation reactivation and associated with reduced lag phases during growth after stress removal. Our findings suggest that mRNA storage within aggresomes confers an advantage for bacterial survival and recovery from stress.

## Main

Bacteria employ liquid-liquid phase separation (LLPS)—a process where biomolecules demix from the cytosol to form concentrated, membraneless condensates—to survive environmental stress^1–4^. While LLPS-driven compartments are well-characterized in eukaryotes, their roles in prokaryotes remain unclear. Work in *Caulobacter crescentus* revealed bacterial ribonucleoprotein particle bodies (BR bodies), RNase E-enriched condensates that balance mRNA decay and storage through dynamic phase transitions^5–7^. Similarly, HP bodies— scaffolded by polyphosphate and the RNA chaperone Hfq—were shown to selectively stabilize translation-related transcripts in diverse bacteria. In *Escherichia coli*, our previous studies identified that ATP-depletion triggers formation of protein-rich aggresomes that sequester stalled translation complexes^9–11^, yet their function as canonical stress granules and mechanisms for mRNA protection remain unresolved.

Here, using *E. coli* as a model system, we demonstrate that aggresomes selectively recruit mRNA via length-dependent partitioning, then protect it through electrostatic exclusion of negatively charged ribonucleases. This mechanism maintains mRNA integrity during stress and enables rapid translation reactivation upon stress relief, ultimately enhancing cellular fitness. Crucially, aggresomes represent a distinct functional paradigm: while condensates like BR bodies coordinate mRNA *degradation and storage*, aggresomes specialize in *preservation primarily* through physical exclusion mechanisms.

## Results

### Aggresomes are enriched with mRNA

ATP depletion is a crucial factor inducing aggresome formation^9,10^. Arsenite is traditionally known as an oxidative stressor causing ATP decrease in eukaryotic cells^12^. Our experiments also demonstrate its effectiveness in inducing cellular ATP depletion in bacterial cells (Extended Data Fig. 1a). By exposing *Escherichia coli* (*E. coli*) cells to 2 mM arsenite, we were able to consistently induce aggresome formation within 30 minutes, as visualized by distinct foci of green fluorescent protein (GFP) fused to the aggresome biomarker HslU (HslU-GFP) in cells (Fig. 1a). To investigate whether RNA also localizes with proteins in bacterial aggresomes, we purified aggresomes from the lysed cells using immunoprecipitation with an antibody specific to HslU-GFP (Fig. 1b). Purified aggresomes exhibited a generally spherical shape and remained intact throughout the process of lysis and purification (Fig. 1c).

**Fig. 1.**
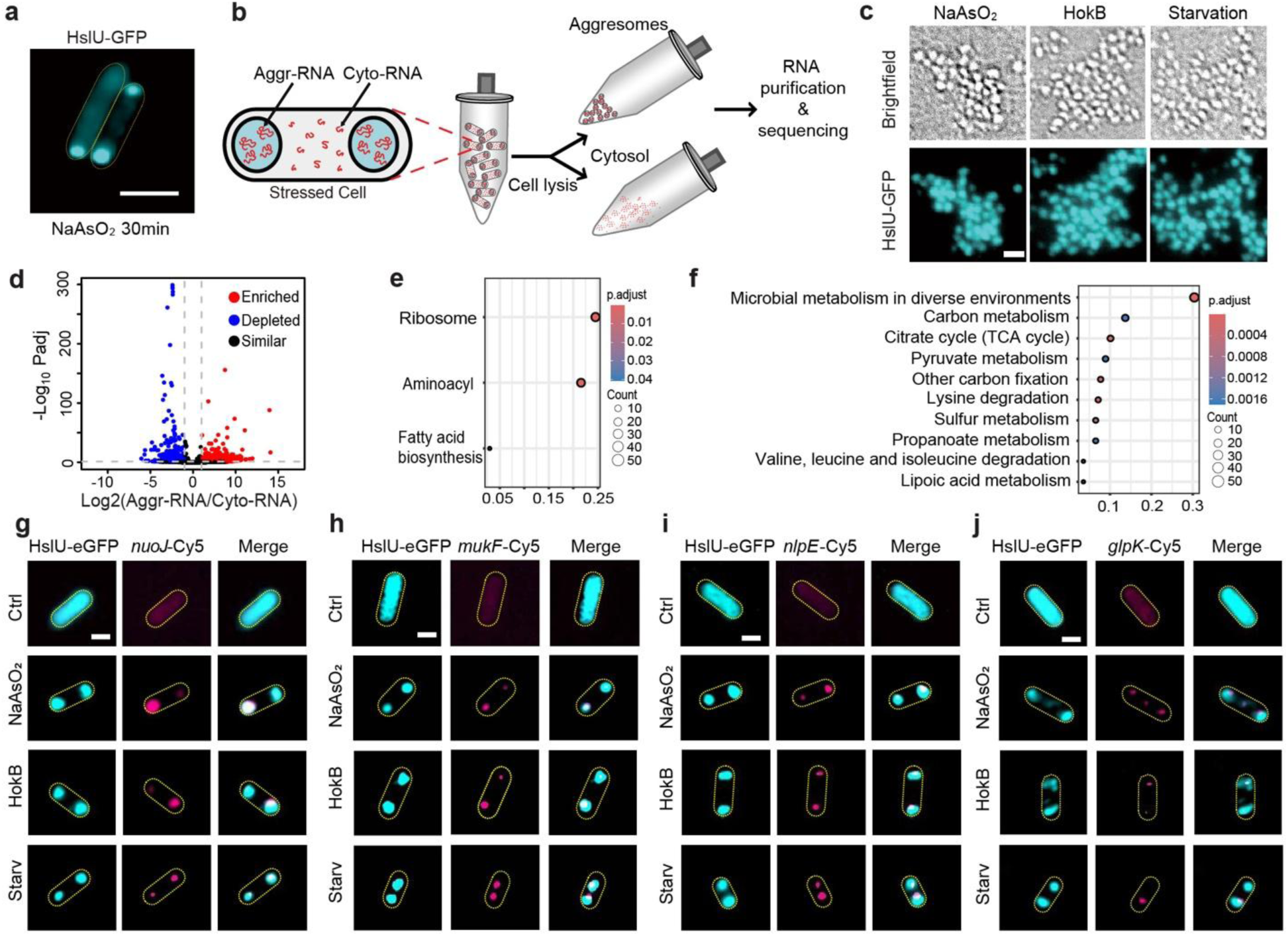
| Aggresome formation requires mRNA. **a**. Formation of distinct HslU-GFP foci (cyan) in cells after 30min of 2mM NaAsO_2_ exposure, cell body indicated (yellow dash), scale bar: 2μm. **b**. Schematic summary of aggresome isolation process for RNA-seq, Aggr-RNA, aggresome RNA; Cyto-RNA, cytosol RNA. **c**. Purified aggresomes from *E. coli* cells treated with 2mM arsenite (NaAsO_2_), HokB overexpression (HokB) and starvation, respectively, scale bar: 1 μm. **d.** Volcano plot of differential RNA abundance between aggresome-enriched fractions (Aggr-RNA) and aggresome-depleted cytosol (Cyto-RNA) (DESeq2 analysis). Significantly enriched RNAs in aggresomes (red; log₂[FC] > 1, adjusted p < 0.01) and depleted RNAs (blue; log₂[FC] < -1, adjusted p < 0.01) are highlighted. Gray dots: non-significant RNAs (|log₂[FC]| ≤ 1 or adjusted p ≥ 0.01). Fold changes represent Aggr-RNA/Cyto-RNA ratios. **e, f**. KEGG pathway analysis of Aggr-RNA (**e**) and Cyto-RNA (**f**). Pathway enrichment analyses a two-sided Fisher’s exact test with Benjamini-Hochberg FDR correction (significant: FDR < 0.05, n = 3 independent biological replicates). **g-j**.Distribution of the aggresome protein HslU, labeled with eGFP, in conjunction with RNA transcripts *nuoJ* (**g**), *mukF* (**h**), *nlpE* (**i**), and *glpK* (**j**), detected using RNA-FISH probes (*nuoJ-Cy5, mukF-Cy5, nlpE-Cy5,* and *glpK-Cy5*). The aggresomes were induced with 2mM arsenite treatment for 30 min (NaAsO_2_), HokB induction for 30 min (HokB), or 24 hours of nutrient starvation (Starv). Control (Ctrl) represents exponentially growing cells without any treatment. Scale bar: 1μm.

Our aggresome extraction protocol preserves RNA integrity, as confirmed by ScreenTape analysis (Extended Data Fig. 1b). Transcriptome analysis of aggresome RNA (Aggr-RNA) and RNA which remained in the cytosol (Cyto-RNA), followed by pairwise correlation analysis (Extended Data Fig. 1c) and differential gene expression analysis (Fig. 1d and Extended Data Fig. 1d, e), revealed that Aggr-RNA was distinct from Cyto-RNA. Kyoto Encyclopedia of Genes and Genomes (KEGG) pathway analysis revealed that aggresomes were significantly enriched in mRNA from genes with functions associated with ribosome, aminoacyl and fatty acid biosynthesis (Fig. 1e). In contrast, mRNA from genes with functions associated with stress response pathways, including microbial metabolism in diverse environments and amino acid degradation, were notably enriched in the cytosol (Fig. 1f). These findings, along with the observation of aggresome formation through phase separation^13^ and the identification of proteins and RNAs associated with stalled translation initiation complexes in aggresomes (Supplementary Table 1), suggest that bacterial aggresomes exhibit similar characteristics to eukaryotic stress granules (SGs).

To validate the transcriptomics data and confirm localisation of specific mRNAs in aggresomes we analysed specific transcripts with significantly different expression levels between aggresomes and cytosol following arsenite induction in the RNAseq data: aggresome-enriched transcripts, *nuoJ*, *nlpE*, *glpK* and *mukF*; and cytosol enriched transcripts *dps, gmhA* and *gppA*. Localisation of aggresome enriched *nuoJ*, *nlpE*, *glpK* and *mukF* was confirmed with RNA fluorescence in situ hybridization (RNA-FISH) (Fig. 1g-j) and real-time polymerase chain reaction (qPCR) analysis (Extended Data Fig. 1f). Both groups of transcripts were labelled with Pepper RNA aptamer^14^ in a background strain genomically expressing HslU-mCherry and visualized with HBC530 green dye under time-resolved structured illumination microscopy (SIM) (Extended Data Fig. 2a-c). In the absence of arsenite, no HslU-mCherry foci formation was observed, and all transcripts were uniformly distributed throughout the cytosol (Extended Data Fig. 2d, Supplementary Video 1-4). After arsenite treatment, *nuoJ, nlpE, glpK* and *mukF* transcripts gradually formed distinct foci, which colocalized with HslU-mCherry foci (Fig. 2a-d; Supplementary Video 5-8), with the rate of foci formation variable across different transcripts (Fig. 2e-h). In contrast, transcripts of *dps*, *gmhA* and *gppA* remained distributed throughout the cytosol after arsenite treatment (Extended Data Fig. 2e; Supplementary Video 9-11). We also tested additional stresses known to lead to cellular ATP depletion: HokB^15^ induction also induced foci formation for *nuoJ, nlpE, glpK* and *mukF* (Extended Data Fig. 2f-i). These results show accumulation of specific transcripts in aggresomes.

**Fig. 2.**
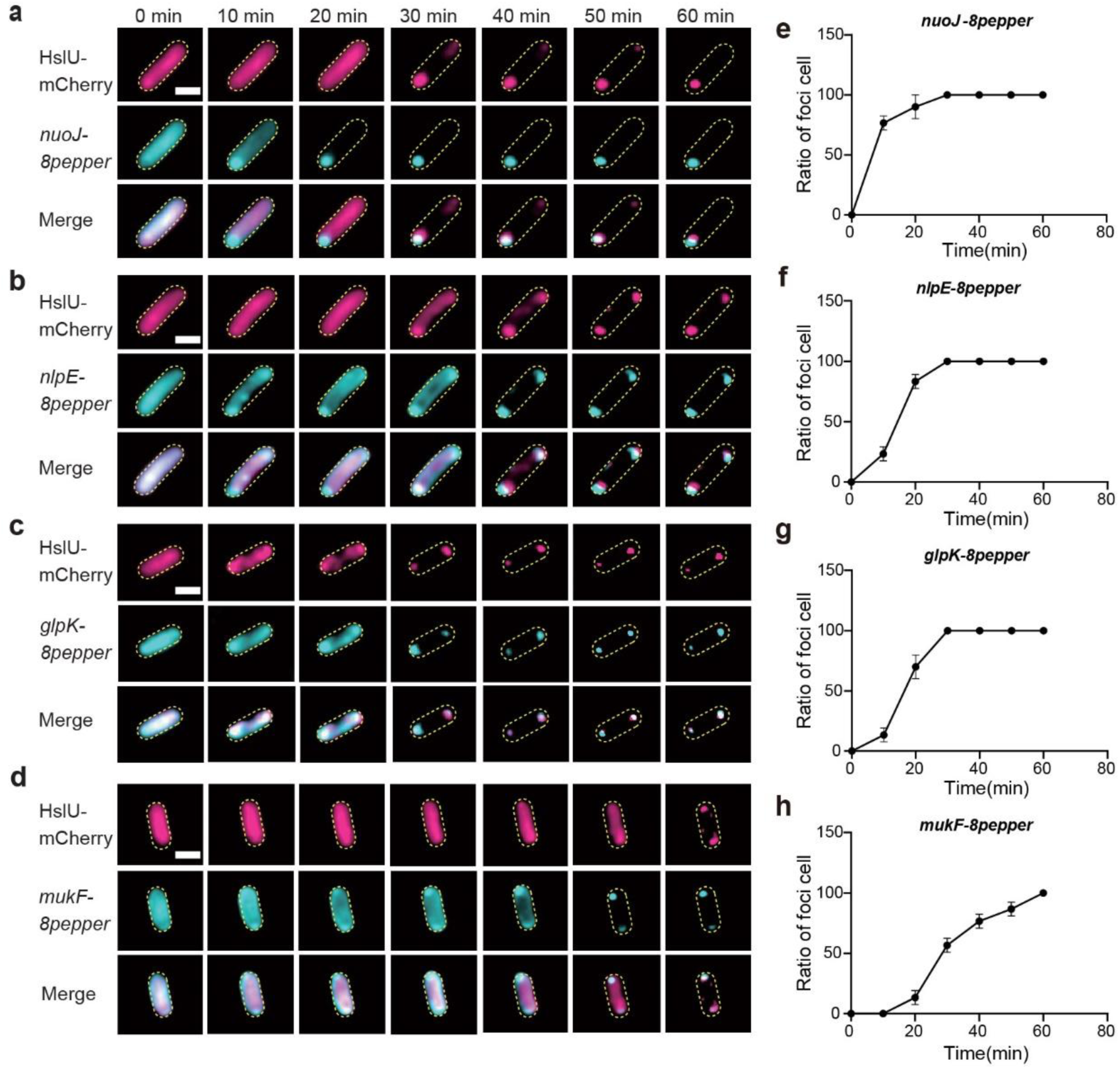
**| Dynamic mRNA partitioning into aggresomes during arsenite stress**. **a-d.** Representative time-lapse images showing aggresomal partitioning of *nuoJ* (**a**), *nlpE* (**b**), *glpK* (**c**), and *mukF* (**d**) mRNAs under sodium arsenite stress (2 mM). Images show dual-color microscopy of HslU-mCherry (magenta; aggresome marker) and mRNA-8pepper labeled by HBC530 (cyan). Scale bar: 1 μm (applies to a-d). **e-h.** Quantitative analysis of cells containing mRNA-containing aggresomes over time for *nuoJ* (**e**), *nlpE* (**f**), *glpK* (**g**), and *mukF* (**h**). Data represent proportion of cells with fluorescent foci (n = 10, error bar indicates SE).

### Prolonged stress triggers aggresome compaction

Quantitative transcriptome analysis, using operonic transcripts as a reference, revealed that mRNA molecules enriched in aggresomes have an average length that exceeds five times the length of those in cytosol (Extended Data Fig. 3a, b). High sensitivity RNA ScreenTape analysis validated these findings with arsenite induction (Fig. 3a-c). Similar results were also observed under stress of HokB induction or starvation (Fig. 3b, c).

**Fig. 3.**
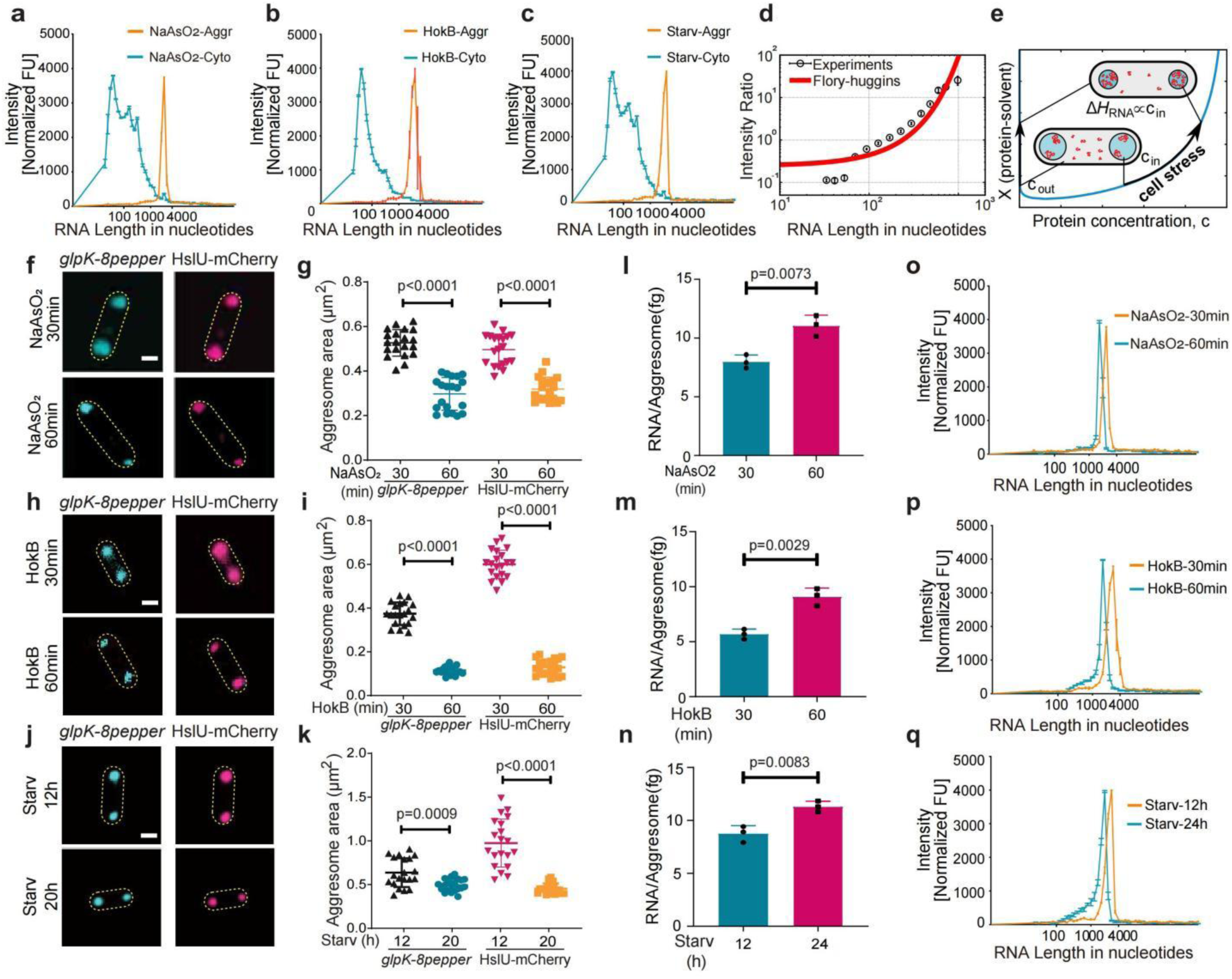
**| Prolonged stress triggers aggresome compaction. a-c**. ScreenTape RNA analysis of Aggresome-RNA (Aggr-RNA) and Cytosol-RNA (Cyto-RNA) from the cells with (**a**) arsenite (NaAsO_2,_ 2 mM, 30 min) treatment, (**b**) HokB induction for 30 min, and (**c**) starvation (Starv) induction for 24 hours (n=3 independent biological replicates; mean ± SE). **d**. Aggresome/Cytoplasm RNA intensity ratio versus RNA length (SD error bars), with overlaid fit (red) based on Flory-Huggin’s theory with optimized parameters Prefac = 0.32 ± 0.08 and ΔH =0.0055 ± 0.0007 k_B_T, goodness-of-fit R_2_=0.989378. **e**. The Flory-Huggins phase diagram for the solvent and protein predicts droplet compaction. The blue curves represent the binodal concentrations in the cytoplasm and in the aggresome. An increasing protein-solvent interaction parameter, χ, (interpreted to correlate with cell stress) leads to an increasing binodal protein concentration inside the aggresome; due to mass conservation this causes the droplets to shrink. **f, h, j**. Representative SIM images of cells with different duration of NaAsO_2_ (2 mM) treatment (**f**), HokB induction (**h**) and starvation (**j**), *glpK-8pepper* represents *glpK-8pepper* RNA stained with HBC530 green dye, HslU-mCherry represents protein HslU labeled by fluorescent protein mCherry. **g, i, k**. Statistical analysis of the area of aggresomes projected onto the camera detector in cells with different duration of NaAsO_2_ (2 mM) treatment (**g**), HokB induction (**i**) and starvation (**k**), based on SIM images (n=20 cells per condition, median ± SE). **l-n**. The mass of RNA per aggresome in cells with different duration of NaAsO_2_ (2 mM) treatment (**l**), HokB induction (**m**) and starvation (**n**), the total mass of RNA in bulk aggresomes (RNA_total_) is measured by a Qubit fluorometer, the number of aggresomes (N_aggresome_) of a given volume was determined by FACS, and RNA/Aggresome is determined by RNA_total_/N_aggresome_. **o-q**. ScreenTape RNA analysis of RNA length in aggresomes from cells with different duration of NaAsO_2_ (2 mM) treatment (**o**), HokB induction (**p**) and starvation (**q**), n=3 independent biological replicates; mean ± SE for **l-q**. Scale bar: 500 nm. Two-sided unpaired Student’s t-test used in comparison; error bars indicate SE.

The association between increased mRNA length and localization within aggresomes aligns with recent studies on LLPS droplets of transcriptional machinery in mammalian cells, where stress granule-enriched mRNAs are often longer than the average length^16^. To understand *why* longer mRNAs preferentially accumulate in aggresomes during stress, we hypothesized that mRNA partitioning follows fundamental biophysical principles: longer RNA chains might gain sufficient favorable interactions (enthalpy) with aggresome proteins to overcome the energy penalty of entering a confined space (entropy). To test this, we applied Flory-Huggins (FH) theory^17^ – a thermodynamic model for polymer solutions – aiming to quantify the energy balance driving mRNA recruitment and explain how RNA length influences partitioning. The model mathematically represents three interacting components: aggresome proteins, RNA, and water, with parameters capturing their pairwise affinities (Methods Modeling and Supplementary Information Fig. 2). We first considered the regime of zero or very low aggresomal RNA concentrations in which the formation and compaction of droplets has been driven by the protein-water interaction parameter. In this regime, the model assumes that RNA has no direct influence on the phase separation process itself. In the steady-state regime of the model, the partitioning of RNA (Fig. 3d, e) is simply driven by an RNA-protein interaction that is favorable compared to the RNA-water interaction, which is enhanced by high protein concentrations in compact droplets. Here, the extent of RNA partitioning only depends on a single emergent parameter, which is modelled as a consensus enthalpy parameter per nucleotide base, Δ*H* (see Equation 11 in Supplementary Information). Therefore, the total enthalpy per RNA molecule is expected to simply increase linearly with its length due to an increased number of nucleotide-protein interactions. We find a reasonable fit to the experimental data spanning two orders of magnitude for the ScreenTape intensity ratio and RNA length using an optimized enthalpy parameter of Δ*H =* (5.5 ±0.7) x 10^−3^ *k*_B_*T* per nucleotide base (Fig. 3d, e). This value quantitatively agrees with our experimental data, providing a comprehensive understanding of the distribution of RNA lengths inside an aggresome at steady-state, through the lens of statistical thermodynamics.

Our model, based on a mass balance analysis of a closed system, simulates the effects of prolonged stress on aggresome dynamics, which indicates that prolonged stress leads to decreased compatibility between proteins and solvents, consequently prompting the compaction of aggresomes. As aggresome compaction sets in, the thermal energy scale increases, thereby resulting in an increased affinity for RNA molecules (Fig. 3e, and Equations 10,11 in Supplementary Information). Notably, the increased affinity for RNA molecules during aggresome compaction is expected from the model to facilitate increased transfer of cytosolic mRNA to the aggresomes.

To experimentally test these model outcomes, we extended arsenite treatment from 30 to 60 minutes. SIM and transmission electron microscopy (TEM) analysis of the aggresome area confirmed the model’s prediction, showing an approximately 50% decrease in the measured cross-sectional area of aggresomes with prolonged stress (Extended Data Fig. 3c, d). This reduction in size was observed for both the aggresome biomarker protein HslU and the Pepper-labeled aggresomal mRNA (Fig. 3f-k; Extended Data Fig. 3e-j). Furthermore, to quantify the mass amount of aggresomal RNA present, we used flow cytometry to determine the absolute number of aggresomes in a given volume and subsequently performed RNA purification. The results show that prolonged arsenite stress leads to an approximate 40% increase in the mass of RNA per aggresome (Fig. 3l) and an approximate 140% increase in the mass of protein per aggresome (Extended Data Fig. 3k). Similar findings were obtained upon aggresome induction with HokB expression and starvation (Fig. 3m, n). These experimental results validate the steady-state model output that prolonged stress triggers aggresome compaction, characterized by a reduction in size concurrent with an increase in mass.

### Dynamic flux of RNA between aggresomes and cytosol

Meanwhile, we observed a time-dependent decrease in mean mRNA length specifically within aggresomes (Fig. 3o-p): shifting from 2,850 ± 210 nt after 30 min arsenite treatment to 2,337 ± 190 nt after 60 min arsenite treatment (*18% decrease*, *P*<0.01; Fig. 4a, b). This *intra-aggresome shortening* contrasts sharply with cytosolic mRNA, which underwent more severe fragmentation (45% length reduction; Fig. 4c, d). We hypothesized this difference reflects compartment-specific protection: while cytosolic RNA undergoes enzymatic degradation, aggresomes provide a sheltered environment that preserves RNA integrity.

**Fig. 4.**
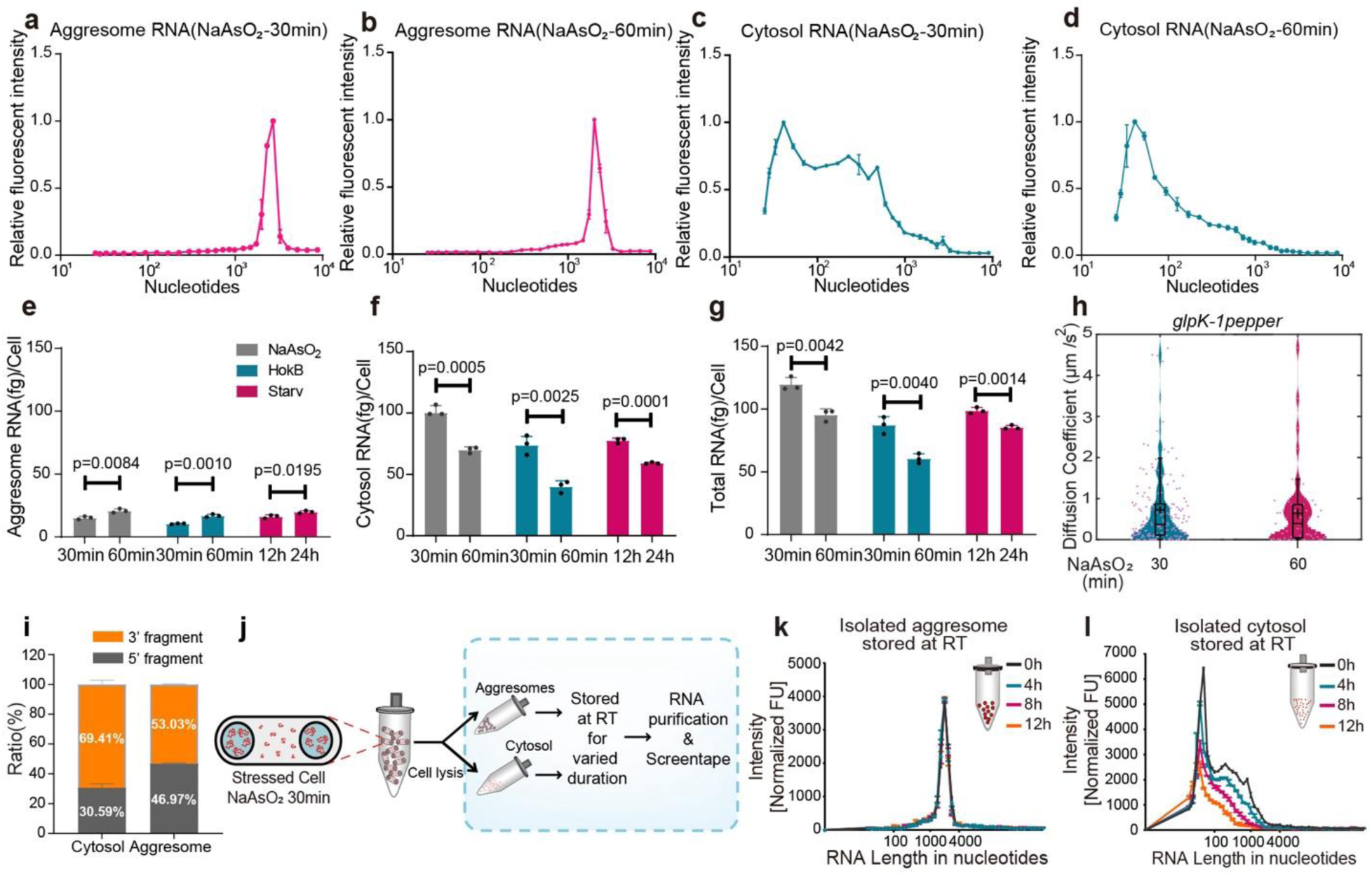
**| RNA is stabilized in aggresomes but degraded in the cytosol under cellular stress. a-d**. ScreenTape RNA analysis of Aggresome-RNA (Aggr-RNA) or Cytosol-RNA (Cyto-RNA) from the cells with arsenite treatment (2mM) for 30 min or 60 min. The fluorescence intensity was normalized using the Min-Max scaling method to standardize the data across samples. **e-g.** Direct measurement of Aggresome RNA/cell, Cytosolic RNA/cell, and Total RNA/cell. **e**. The mass of RNA in aggresomes per cell subjected to different stresses for varying durations. **f**. The mass of RNA in cytosol per cell subjected to different stresses for varying durations. **g**. The mass of total RNA per cell subjected to different stresses for varying durations. **h**. Distribution of diffusion coefficients as violin jitter plot for *glpK* transcript in aggresomes with arsenite exposure for 30 minutes or 60 minutes. **i**. Distribution of RNA-seq reads mapping to the 3’ and 5’ fragments of operons in the cytosol and aggresomes. **j-l**. Differential RNA stability in aggresomes and cytosol *in vitro*. **j**. Schematic summary of *in vitro* analysis. **k**. Isolated aggresomes protecting RNA *in vitro*. RNAs were extracted from isolated aggresomes that have been stored at room temperature for 0, 4, 8, and 12 hours, respectively. Isolated aggresomes derived from cells treated with arsenite for 30 minutes. **l**. RNA undergoing degradation in isolated cytosol *in vitro*. RNAs were extracted from isolated cytosol that have been stored at room temperature for 0, 4, 8, and 12 hours, respectively. Isolated cytosol derived from cells treated with arsenite for 30 minutes. **a-g, i, k, l**: n=3 independent biological replicates, mean ± SE. Two-sided unpaired Student’s t-test used in comparison; error bars indicate SE.

To account for this effect, we expended the steady-state model to include the potential effects of dynamic RNA mobility outside of the aggresome, and RNA flux between the cytosol and aggresome. Here, the adapted model comprises dynamic coupled processes which involve diffusion of cytosolic RNA, the incorporation of RNA into an aggresome, and then the potential exit of the RNA from the aggresome (for full details see Supplementary Information Fig. 3). In this dynamic framework, although the strength of enthalpic binding of RNA in the aggresome is higher for longer RNA, as incorporated into the steady-state model, the rate of RNA diffusion within the cytosol is slower for longer RNA, described by *K_diff,C_ = K^0^_diff_N _R_ ^-^*^1^*^/^*^2^, where *N_R_* is the RNA length in terms of number of nucleotide bases. However, after an RNA molecule is incorporated into an aggresome it may then exit with a probability which decreases with increasing RNA length since it is proportional to the Boltzmann factor exp(*-N_R_ΔH/k_B_T*). When RNA is outside the aggresome in the cytosol, the dynamic model assumes it may undergo enzymatic degradation, described as a process whereby each bond between nucleotides may break with a rate *K_deg_*. The dynamic interplay between diffusion and degradation then leads to a net depletion of RNA from the aggresome over time. This depletion process begins with shorter RNA chains, as they have a higher probability of exiting the aggresome. Some of these shorter RNA fragments may then diffuse back into the aggresome, creating a dynamic equilibrium. These processes are governed by master equations (see Equations 15 and 16-17 in the Supplementary Information), which describe the time evolution of RNA concentrations in the aggresome and cytosol. These coupled dynamic processes result in a complex interplay between diffusion, degradation, and protection within the aggresome, ultimately shaping the RNA length distribution over time. A clear result of this dynamic RNA flux is that at later times following extended arsentite treatment the dynamic model predicts a decrease in the mean RNA length inside the aggresome, even though all the RNA degradation *per se* has occurred *outside* the aggresome in the cytosol. A further expectation of the model is that there is typically a much greater decrease in the mean RNA length in the cytosol over comparable time scales.

To test these dynamic model outcomes, we quantified the levels of aggresomal, cytosolic and total RNA from cells subjected to arsenite stress, HokB induction, or starvation for a range of durations. These direct measurements revealed that prolonged stress triggers decrease in total cellular RNA, increase in aggresomal RNA and decrease in cytosolic RNA (Fig. 4e-g). This compartment-specific redistribution was consistent across arsenite, HokB induction, and starvation stresses.

To determine how cellular stress affects the mobility of aggresomal mRNA, we used high-sensitivity millisecond single-molecule Slimfield microscopy^18^ to track its spatial localization inside living cells (Supplementary Table 2, Supplementary Video 12, and Extended Data Fig. 3l). By photobleaching the majority of labeled mRNA molecules in live cells, we could track individual molecules and determine their apparent diffusion coefficient from the initial gradient of the mean square displacement relative to tracking time. After 30 minutes incubation with arsenite, the mean diffusion coefficient of the four aggresome mRNA transcripts ranged from 0.43 ± 0.06 µm^2^/s (±SE; number of tracks *N* = 159) for *nuoJ* to 0.84 ± 0.12 µm^2^/s (*N* = 129) for *mukF* (Fig. 4h and Supplementary Table 2). These values were significantly higher than the mean diffusion coefficient of the aggresome protein biomarker HslU (approximately 0.2 µm^2^/s), consistent with single mRNA molecules exhibiting liquid-like diffusion within aggresomes. Prolonged incubation for 60 minutes with arsenite resulted in a mean diffusion coefficient ranging from 0.36 ± 0.06 µm^2^/s (*N* = 147) for *nuoJ* to 0.64 ± 0.13 µm^2^/s (*N* = 49) for *glpK*. These findings suggest a trend towards a more viscous state within the aggresome during prolonged stress, affirming stress-induced compaction characterized by a simultaneous reduction in size and an increase in mass.

### Aggresomes protect mRNA by exclusion of mRNA ribonucleases

Bacterial gene expression is tightly regulated through mRNA degradation. The degradation pathways predominantly involve endonucleolytic cleavage of primary transcripts into 5′ and 3′ fragments. The 5′ fragment, which is unprotected at its 3′ end, is then rapidly degraded by 3′ to 5′ exonucleases^19^. Accordingly, in total RNA populations, 3’ fragments typically accumulate at higher proportions than 5’ fragments due to differential stabilities. RNA-seq reads mapping analysis of aggresome-associated RNA revealed comparable proportions of 3’ and 5’ fragments, contrasting sharply with cytosolic RNA, where 3’ fragments were significantly enriched (Fig. 4i). To further explore whether aggresomes play a role in protecting sequestered mRNA, we extracted aggresomes and cytosols from arsenite stressed cells. Subsequently, these isolated aggresomes and cytosols were stored at room temperature for varying durations (0, 4, 8, and 12 hours). RNA was then purified for ScreenTape analysis (Fig. 4j). We observed preservation of RNA transcriptome within aggresomes over prolonged periods (Fig. 4k) while cytosolic RNA gradually degraded over time (Fig. 4l).

These live-cell and *in vitro* findings together suggest that aggresomes prevent RNA degradation. This was confirmed by qPCR assays to compare the degradation rates of individual mRNA transcripts in aggresomes and cytosol. We analysed *talB* and *gltI* transcripts. The qPCR data showed after prolonged stress, the relative levels of the transcripts were significantly reduced in the cytosol compared to the levels in the aggresome (Extended Data Fig. 4a-d). These observations suggest that RNA accumulation within aggresomes confers protection against cytosolic degradation pathways.

To investigate the mechanism of mRNA protection in aggresomes, we used affinity purification mass spectrometry (AP-MS) to explore proteomes enriched or depleted in aggresomes. AP-MS analysis output in the Search Tool for the Retrieval of Interacting Genes/Proteins (STRING) database revealed distinct protein-protein interaction (PPI) network topologies for aggresomes compared to the cytosol, with mean clustering coefficients of 0.673 and 0.496, respectively (Extended Data Fig. 4e-g). Of particular significance, ribonucleases specialized for rRNA processing (RNG and RIA) and for tRNA processing (RnpA) were enriched in aggresomes, whereas RNAses responsible for mRNA degradation, including Exoribonuclease II (RNB), Ribonuclease BN (RBN) and Oligoribonuclease (ORN), were depleted from aggresomes (Fig. 5a). SIM imaging using the HslU-GFP aggresome biomarker confirmed that mCherry-fused RNB, RBN and ORN under the control of their native promoters, were excluded from aggresomes induced by arsenite treatment (Fig. 5b-d). Similar results were observed under stress of HokB induction or starvation (Extended Data Fig. 4h-m). These data suggest that mRNA ribonucleases might be excluded from aggresomes.

**Fig. 5.**
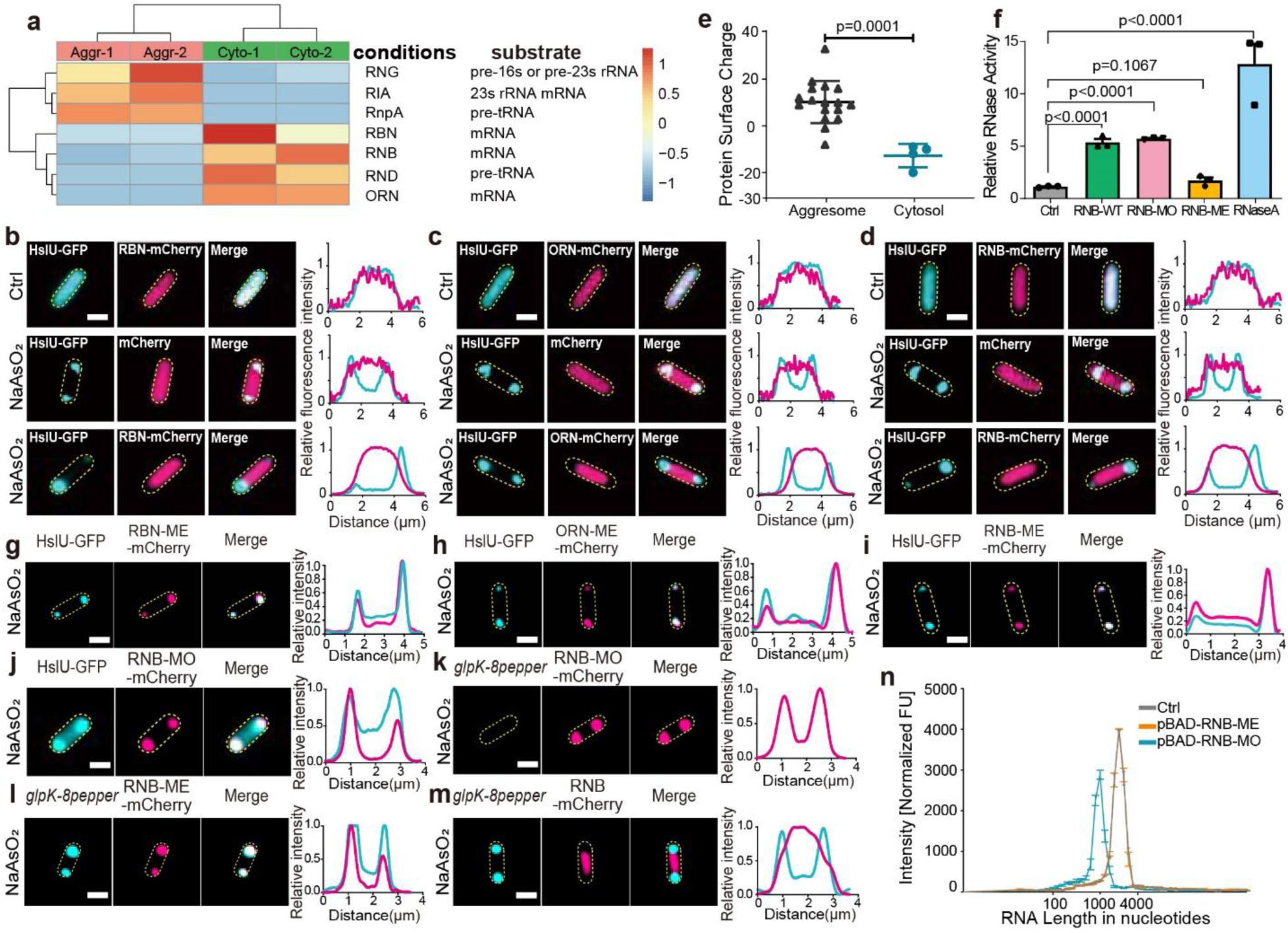
| Aggresome protects mRNA by excluding ribonucleases. **a**. Heatmap of RNase abundance in aggresomes (Aggr) vs. cytosol (Cyto) (log₂-normalized). **b-d**. Distribution of ribonucleases RBN (**b**), ORN (**c**), RNB (**d**) in cells with chromosomally labeled HslU-EGFP under arsenite treatment. Each ribonuclease is labeled by mCherry. **e**. Statistical analysis of protein surface charges of RNA binding proteins in aggresomes or that of RNases in cytosol (n=3 independent biological replicates, mean ± SE). **f**. RNase enzyme activity measurement. RNase A serves as the positive control, while "Ctrl" denotes the buffer-only control (n=3 independent biological replicates, mean ± SE). **g-j**. Distribution of ribonuclease mutants RBN-ME (**g**), ORN-ME (**h**), RNB-ME (**i**), RNB-MO (**j**) in cells with chromosomally labeled HslU-EGFP under arsenite treatment. Each mutant protein is labeled by mCherry. **k-m**. Localization of RNB variants in glpK-8pepper/HBC530 cells: RNB-MO (**k**), RNB-ME (**l**), wild-type RNB (**m**) with intensity profiles. **n**. ScreenTape analysis of aggresome RNA from cells overexpression RNB-ME or RNB-MO after 30 min of 2 mM arsenite treatment. Ctrl, aggresome RNA from wild-type cells after 30 min of 2 mM arsenite treatment (n=3 independent biological replicates, mean ± SE). ME: Full enzymatic center mutation; MO: Mutation outside RNA-binding/catalytic domains. Two-sided unpaired Student’s t-test used in comparison; error bars indicate SE.

### Protein surface charge excludes mRNA RNases from aggresomes

Our statistical thermodynamics steady-state model, based on Florey-Huggins theory, provided an explanation for the exclusion of RNases from aggresome droplets. In this model, we incorporated different interaction parameters to account for the higher affinity of ribonucleases to the water solvent and their repulsion away from the aggresome. Using bioinformatics analysis, we revealed a significant net negative surface charge that distinguishes RNA nucleases (RNB, -20; RBN, -9; ORN, -10) from most RNA binding proteins (RBPs) present within aggresomes (Fig. 5e, f and Extended Data Fig. 5a). Building on these findings, we hypothesized that the exclusion of ribonucleases from aggresomes is influenced by negative charge repulsion, possibly from the RNA molecules, which have a high net negative charge due to the phosphate backbone, already present within the condensate.

To test this hypothesis, we conducted mutagenesis by substituting all aspartic acid (D) or glutamic acid (E) residues across the entire enzyme center with alanine (A), which resulted in a drastic change in surface charge (RNB-ME, 4; RBN-ME, 26; ORN-ME, 20) (Extended Data Fig.5b) while also disrupting the RNA binding motif and catalytic centre. In this scenario, the mutated ribonucleases were able to enter aggresomes without influencing aggresome RNA condensation. SIM imaging following arsenite induction confirmed colocalization in distinct foci of mCherry-fused RNB-ME, RBN-ME, or ORN-ME, with HslU-GFP foci (Fig. 3h-j). These results suggest that surface charge leads to exclusion of mRNA ribonucleases from aggresomes. Furthermore, RNB plays a significant role in mRNA degradation, which enzymatically hydrolyzes single-stranded polyribonucleotides in a processive 3’ to 5’ direction^20^. We conducted mutagenesis by substituting only the D and E residues located outside the RNA binding motif and catalytic center of RNB with an A residue (RNB-MO). This resulted in a moderate positive surface charge (Extended Data Fig. 5c), while the protein’s enzymatic activity remained intact (Fig. 5d). SIM imaging following arsenite induction also showed entry of positively charged RNB-MO into aggresomes and depletion of RNA (Fig. 5k-n, Extended Data Fig. 5d, e). RNA ScreenTape analysis confirmed these results (Fig. 5o). These data demonstrate that negative protein surface charge mediates electrostatic exclusion of ribonucleases (RNases) from aggresomes, thereby protecting sequestered mRNA from degradation.

### Aggresome formation promotes recovery after stress

Time-resolved SIM showed that, following removal of arsenite stress, the transcripts of *glpK* (Fig. 6a) and *nuoJ, nlpE, mukF* (Extended Data Fig. 6a-d) were released from aggresomes into the cytosol over a period of 1-3 hours. Previous mass spectrometry and RNA-seq analysis showed that both mRNA and protein products of *glpK* accumulated in aggresomes at high levels compared to the cytosol during stress (Extended Data Fig. 6e, f). To assess whether released mRNA is translated, we used an arabinose-inducible fluorescent reporter of the GlpK protein. Following arsenite induction, GlpK-GFP localized to aggresomes as expected. We then photobleached the entire cell and removed arsenite from the imaging buffer. To prevent the influence of newly transcribed mRNA, we introduced sub-Minimum Inhibitory Concentration (sub-MIC) rifampicin to inhibit RNA synthesis. Additionally, to account for potential pre-existing fluorescent proteins maturing after photobleaching, we supplemented the medium with tetracycline, which inhibits bacterial protein synthesis by targeting the 30S ribosomal subunit. Consequently, in cells treated with rifampicin alone, the reappearance of GlpK-GFP fluorescence can be attributed to both new synthesis of the protein from aggresome-released RNA and pre-existing fluorescent proteins maturing after photobleaching. In contrast, in cells treated with both rifampicin and tetracycline, the reappearance of GlpK-GFP fluorescence can be attributed solely to pre-existing fluorescent proteins maturing after photobleaching. Importantly, the reappearance of the fluorescence in cells treated with rifampicin alone is significantly higher than that in cells treated with both rifampicin and tetracycline (Fig. 6b, c), thereby confirming the release of preserved mRNA from aggresomes for subsequent protein synthesis.

**Fig. 6.**
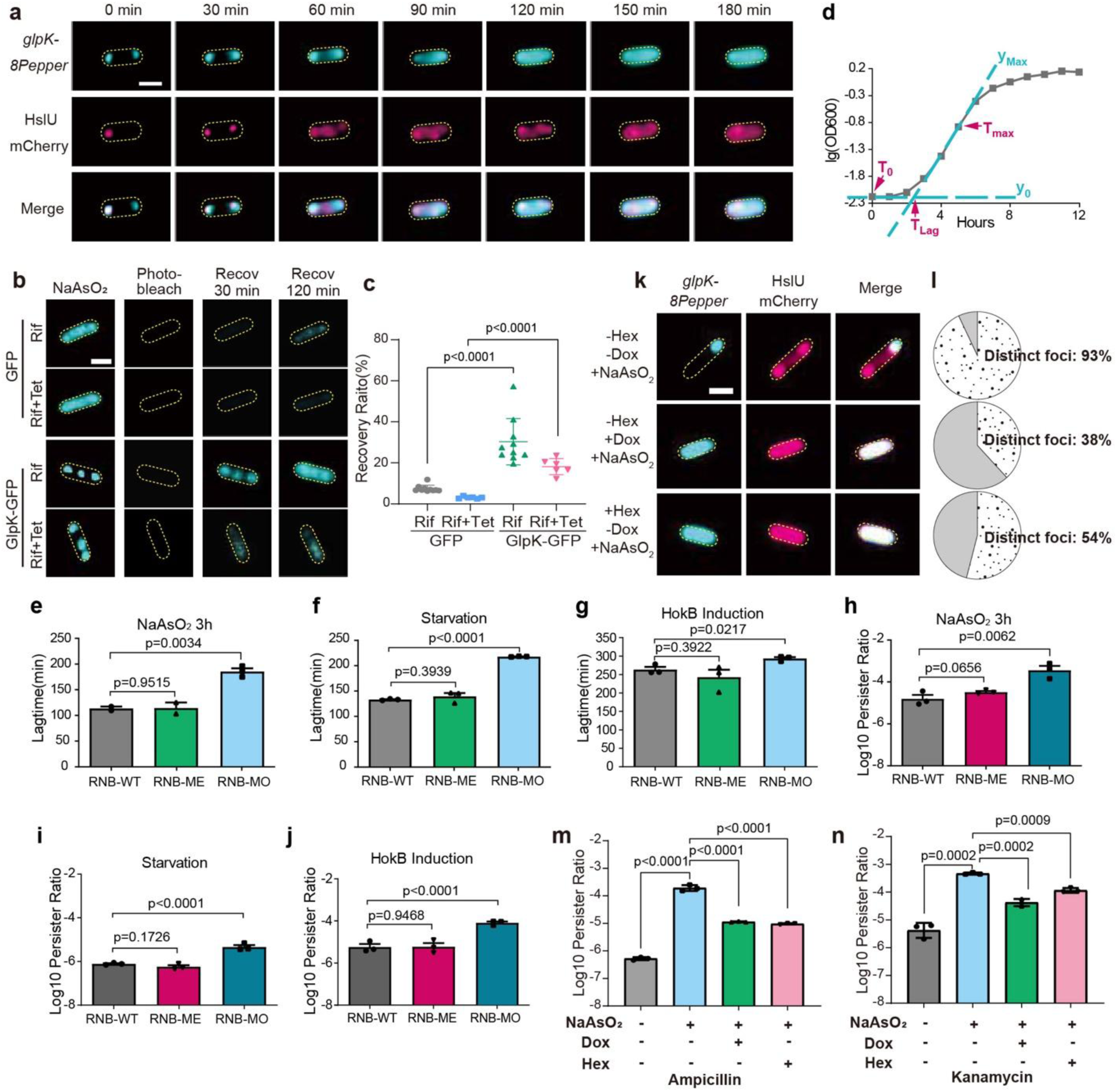
| The translation of released mRNA from aggresomes promotes cell fitness. **a**. Releasing of *glpK-8pepper* mRNA from aggresomes into cytoplasm upon arsenite removal. **b**. Translation of *gfp* or *glpK-gfp* mRNA after aggresome disassembly. Rif, rifampicin; Tet, tetracycline; Recov, recovery time. **c**. Statistical analysis of GFP fluorescence recovery ratio at 120 min after photobleaching, depicted from (**b**) (n=10 tracked cells, mean ± SE). **d**. Lag time calculation method (tangent intersection at T₀ and T_max_). **e-g**. Lag time of strains containing wild-type RNB or RNB-MO and RNB-ME after exposure to 2 mM arsenite for 3 hours (**e**), starvation for 24 hours (**f**) or HokB expression with 0.001% arabinose (w/v) for 30 minutes (**g**). **h-j**. Persister cell ratio in strains containing wild-type RNB or the mutant RNB-MO and RNB-ME after exposure to 2 mM arsenite for 3 hours (**h**), starvation for 24 hours (**i**) or HokB expression with 0.001% arabinose (w/v) for 30 minutes (**j**). **k**. Representative fluorescence images of cells with different chemical combination treatment. Ars, arsenite; Dox, doxorubicin; Hex, 1,6-hexanediol. **l.** Statistical analysis of cells with distinct mRNA foci after different chemical combination treatment, data from (**k**) (n=100 cells). **m, n**. Cell survival rate (log scale) after 4 hours (**m**) ampicillin (**n**) kanamycin killing. **e-n:** n=3 independent biological replicates, mean ± SE. Scale bar: 1 μm. Two-sided unpaired Student’s t-test used in comparison; error bars indicate SE.

Given that preserved mRNA could be translated, we hypothesized that mRNA protection within aggresomes might enable a faster response and reduced lag times following alleviation of stress. To test this, we used RNB-MO mutants to degrade aggresomal mRNA. Strains carrying the mutant RNB-MO exhibited longer lag times following stress exposure compared to those harboring the wild-type RNB (Fig. 6d-g). Interestingly, this prolonged lag time correlated with an increase in the overall fraction of persister cells within the populations (Fig. 6h-j). These observations suggest a potential link between RNA availability and cellular recovery kinetics.

To determine if mRNA protection by aggresomes confers survival advantages, we performed screening of small chemical compounds (Fig. 6k; Extended Data Fig. 7), identifying conditions of media supplementation with 1,6-hexanediol (Hex) or doxorubicin (Dox). Hex is commonly used to study biomolecular LLPS, since it dissolves liquid condensates but not solid-like aggregates via the disruption of RNA-protein interactions^21^, while Dox is a nucleic acid intercalator which inhibits RNA gelation *in vitro* and dissolves RNA nuclear foci in eukaryotic cells^22^ . Hex or Dox alone had no effects on inducing aggresome formation or promoting persister formation (Extended Data Fig. 8). However, aggresome formation was impaired in cells treated with Hex or Dox after arsenite exposure (Fig. 6k, l). We compared the survival difference between untreated cells, and those treated with Hex or Dox during arsenite induction or other cell stress factors. Untreated cells, which exhibited expected aggresome formation, consistently showed a significantly higher persister ratio after different types of antibiotic treatment. On the contrary, cells treated with Hex or Dox showed significantly reduced aggresome formation correlated to a lower prevalence of persisters (Fig. 6m, n). Taken together, these observations indicate that aggresomes enable protection of captured mRNA from ribonuclease degradation, which facilitates translation of released mRNA following stress removal, resulting in significantly increased cell fitness during stress conditions.

## Discussion

Our study establishes *E. coli* aggresomes as bona fide bacterial stress granules that preserve mRNA through a unique mechanism: electrostatic exclusion of ribonucleases. This distinguishes aggresomes fundamentally from other bacterial condensates. For example, BR bodies in *Caulobacter crescentus* incorporate RNase E to balance mRNA decay and storage^5,7,23^, whereas aggresomes actively repel RNases via charge-based exclusion (Fig. 5). This mechanistic divergence defines a critical functional distinction—mRNA preservation in aggresomes versus regulated degradation in BR bodies. Further investigations reveal a hierarchical mRNA protection strategy within aggresomes, beginning with the primary electrostatic repulsion that ensures mRNA-degrading RNases are effectively excluded. This crucial initial defense is corroborated by charge-reversal mutagenesis (RNB-MO), which, by altering the charge characteristics, allows RNase entry and consequently triggers mRNA degradation, thereby directly affirming the importance of electrostatic repulsion in safeguarding mRNA. Beyond this key electrostatic shield, an enrichment of non-coding RNAs, such as *ssrA* and *csrB*, within aggresomes suggests an additional layer of stabilization, potentially through mechanisms like ribosome rescue and interference with mRNA turnover pathways. Future work is needed to elucidate the potential involvement of RNA chaperones such as Hfq or ProQ in this process.

Using experimental data and polymer physics modeling based on FH theory, we elucidate aggresome dynamics under stress: Long mRNAs are selectively sequestered first, consistent with length-dependent partitioning driven by enthalpic gains. As stress persists, aggresomes compact, progressively recruiting shorter transcripts to achieve higher density and mass (Fig. 3f–q), but cytosolic RNA is degraded (Fig. 4). While our polymer physics model indicate qualitative trends (e.g., phase-separation dependencies), its quantitative predictions require cautious interpretation due to inherent limitations, such as the assumption of phantom chains, the potential overestimation of dense-phase volume fractions, and the neglect of local molecular correlations beyond mean-field approximations. Critically, any alternative model must adhere to mass balance constraints: over experimental timescales, RNA nucleotides are conserved (no significant new expression or diffusion). Thus, changes in RNA distribution must account for (i) cell volume expansion/shrinking, (ii) RNA accumulation in aggresomes (concentration × volume), and/or (iii) RNA changes outside aggresomes (concentration × cytosolic volume).

The functional implications of mRNA preservation within aggresomes are profound, directly contributing to cellular robustness and adaptability. By safeguarding mRNA integrity during periods of stress, aggresomes enable a rapid and efficient reactivation of translation immediately upon stress removal, significantly reducing the recovery lag time and thereby enhancing overall cellular fitness (Fig. 6). This rapid recovery mechanism is a critical advantage for bacteria thriving in fluctuating environments, allowing them to quickly resume metabolic activities and proliferation. The importance of this protective mechanism is further underscored by experiments disrupting it; for example, through surface charge mutagenesis of RNases, which allows their entry into aggresomes. Such disruptions not only prolong the recovery lags post-stress, directly linking the efficacy of mRNA safeguarding to the cell’s ability to survive otherwise lethal challenges. This functional connection highlights aggresomes as crucial determinants of bacterial resilience.

Despite the advances presented in our study, several important questions remain, opening avenues for future research. A key area for exploration is how specific non-coding RNAs, particularly *ssrA* and *csrB*, fine-tune the protective mechanisms within aggresomes. Understanding their precise roles could reveal novel regulatory layers of mRNA stability. Furthermore, the potential cooperation between aggresomes and other bacterial condensates, such as HP bodies, warrants investigation. Elucidating such interactions could unveil a broader network of cellular organization and stress response mechanisms. Finally, the intriguing observation that overexpression of RNase E can compromise aggresome integrity and formation presents a compelling question for future studies. Unraveling the molecular basis of this disruption could provide critical insights into the delicate balance required for aggresome assembly and function^24^. Collectively, these findings not only refine our understanding of bacterial stress granule functionality but also firmly position electrostatic exclusion as a foundational principle of mRNA preservation, critical for bacterial adaptation and survival in dynamic and challenging environments.

## Methods

### Bacterial strains and plasmids construction

Generation of strains with a chromosomal gene-fluorescent protein translational fusion were performed using λ-red mediated recombination system following previous established protocols^25^. Specifically, DNA fragments encoding GFP or mCherry were cloned into the pSCS3V31C^26^ plasmid using Gibson Assembly (NEB #E2611), replacing the stop cassette between XhoI and BamHI restriction sites. Next, the GFP-Toxin-Cm^R^ or mCherry-Toxin-Cm^R^ fragment was PCR-amplified from modified plasmid using Phanta Super-Fidelity DNA Polymerase (Vazyme #P501) with custom primers containing gene-specific ∼50bp homology arms. Electrocompetent cells expressing λ-Red genes from pSIM6^27^ were transformed with 500 ng PCR product via electroporation (2.5 kV, 5 ms pulse) in 2 mm gap cuvettes. The transformed cells were plated on selection plates containing relative antibiotics. The Toxin-CmR cassette was subsequently removed from the chromosome via another round of λ-red mediated recombination, utilizing a counter selection template. Finally, the cells were plated on counter selection plates containing rhamnose to activate the toxin.

To construct strains with an overexpression plasmid or plasmids with native promoters of target proteins, the bacterial recombinant protein vector pBAD and p15G was used in this study, respectively. The target protein fragment was first amplified from wild type *E. coli* MG1655, recombined with the pBAD or p15G plasmid via homologous recombination and then transformed into electrocompetent DH5α cells. The transformed cells were plated on selection plates containing relative antibiotics. Correct recombinant plasmids were subsequently extracted using the FastPure Plasmid Mini Kit (Vazyme DC201-01) and the plasmids were transformed into appropriate electrocompetent cells according to the experimental requirements. All the plasmids used in this study were listed in Supplementary Table 3 and the primers for strains and plasmids construction were listed in Supplementary Table 4.

### RNA labeling

To perform RNA labeling experiments, an mRNA-pepper recombinant plasmid which involved fusing an 8-pepper tag to the 3’ end of the target mRNA was first constructed. Overnight bacterial cultures expressing pepper-mRNA were diluted by 1:100 into fresh LB and incubated in a shaker (220 rpm) for 2 hr. Next, the expression of the target mRNA was induced by adding 0.001% arabinose (w/v) and incubated in a shaker (220 rpm) for an additional 4 hr. The treated cells were then collected by centrifugation at 4000g for 5 mins and suspended with Imaging Buffer (40 mM HEPES, pH 7.4, 100 mM KCl, 5 mM MgCl_2_ buffer, 1:100 HBC ligands). Cells were stained for 5 mins, protected from light at room temperature.

### Brightfield and fluorescence microscopy

Brightfield and fluorescence imaging were performed on a Nikon Ti2-E inverted fluorescence microscope. Illumination was provided by a solid-state laser at wavelength 488 nm for GFP. The fluorescence emission signal of cells was imaged onto a pco.edge camera.

### SIM microscopy

To perform SIM super-resolution microscopy, overnight bacterial cultures were diluted 1:100 into fresh LB medium and incubated in a shaker at 220 rpm for 2 hr. To induce the expression of the target protein, 0.001% arabinose (w/v) was added, and samples were further incubated in a shaker at 220 rpm for 2 hr. Imaging was carried out using a Nikon N-SIM S microscope, with illumination provided by solid-state lasers at wavelengths of 405 nm for Tag-BFP and Hoechst, 488 nm for GFP and 561 nm for mCherry.

### Time-resolved widefield imaging

The Flow Cell System FCS2 (Bioptechs) system was used to perform the time-resolved widefield imaging^28^. Treated bacterial cultures were harvested by centrifugation, washed, suspended in PBS buffer and then imaged on a gel-pad made up of 3% low gelling temperature agarose, with a cell culture to gel-pad volume ratio of 1:10. The gel-pad was positioned at the center of the FCS2 chamber and surrounded by LB liquid medium buffer containing 20 mM NaAsO_2_. The cells were observed under bright-field/epifluorescence illumination at 37 ℃, with images captured every five mins over a period of 180 mins. Imaging was carried out using a Nikon N-SIM S microscope.

### Aggresome isolation and extraction of aggresome RNA

*Bacterial Culture and Stress Induction*: MG1655 bacterial cultures expressing chromosomal fusion HslU-GFP were grown in LB medium in a shaking incubator at 220 rpm and 37°C until reaching exponential phase (OD600 ≈ 0.4-0.6). For stress induction, three conditions were applied: (1) NaAsO2 Treatment: 2 mM NaAsO2 was added to the culture, incubating for various durations; (2) Starvation Stress: exponential phase cultures were continued in the same nutrient-depleted medium for extended periods; (3) HokB Induction: MG1655 strain with HslU-GFP fusion and pBAD-HokB plasmid was grown to exponential phase, then induced with 0.001% (w/v) arabinose for various durations. For all conditions, samples were collected at predetermined time points for further analysis, with specific durations adjusted according to experimental requirements. The formation of aggresomes was confirmed using fluorescent microscopy. The OD600 of the culture was determined using a spectrophotometer. *Cell Harvesting and Lysis:* Cells were collected by centrifugation at 6,500g for 5 minutes and washed once with PBS buffer. All subsequent procedures were performed at 4°C unless otherwise specified. The cells were resuspended in lysis buffer (50 mM Tris-HCl pH 7.4, 100 mM KOAc, 2 mM MgOAc, 0.5 mM DTT, 50 μg/mL Heparin, 0.1% Triton, complete mini EDTA-free protease inhibitor, and 1 U/μL RNasin Plus RNase Inhibitor, 5mg/mL lysozyme), with the buffer volume adjusted to 1 mL per 10 mL of cells at OD600 = 0.5. Cells were lysed using the repeated freeze (liquid nitrogen) - thaw (4°C) method^24^, ensuring effective disruption of cellular structure. The lysates were centrifuged at 1,000g for 5 minutes at 4°C to remove cell debris. *Aggresome Enrichment:* The remaining 950 μL of supernatant was centrifuged at 18,000g for 20 minutes at 4°C to enrich aggresomes. The resulting pellet was washed twice with 1 mL of lysis buffer. The collected aggresomes were confirmed under a fluorescence microscope to ensure proper isolation and integrity. Optionally, affinity purification using anti-GFP antibody (MBL, 048-3, clone 1E4. 1:100 in dilultion in PBS) could be performed, involving centrifugation, resuspension, rotation, and washing steps. Notably, comparative studies indicated that this optional step does not alter RNA profiles (Extended Data Fig. 15). *RNA Extraction from Aggresomes:* The aggresome pellet was resuspended in 100 μg/mL Proteinase K solution and incubated for 15 minutes at 37°C. RNA was subsequently extracted and purified using the Bacteria RNA Extraction Kit (Vazyme, R403-01), substituting the RNA isolater Total RNA Extraction Reagent with RNAiso Plus reagent (Takara, #9109). The purified RNA was eluted in equal volumes of nuclease-free water, and the integrity of the control samples was validated using ScreenTape analysis (Extended Data Fig. 5c).

### Direct Measurement of Aggresomal, Cytosolic and Total Cellular RNA Content

After stress stimulation, the bacterial culture underwent the following processes to assess RNA content: Cell Density Measurement. Cell density was measured by preparing serial dilutions, plating on appropriate growth media, incubating for colony formation, counting Colony Forming Units (CFU) the next day, and calculating cells per mL to establish a baseline. Next, direct RNA measurement from different cellular fractions was performed by lysing cells according to the above protocol and taking equal volumes of cell lysate for aggresome isolation and RNA purification, cytosol isolation and RNA purification, and total RNA purification. Aggresome and Cytosol isolation. The process was initiated by centrifuging the cell lysate at low speed (at 1000 g for 5 mins at 4℃) to remove debris, carefully collecting the supernatant; this supernatant was then subjected to high-speed centrifugation (at 18,000g for 20 mins at 4 ℃) to separate aggresomes from the cytosolic fraction. Cytosolic fraction in the supernatant is carefully collected. The pellet, which composed of aggresomes, was washed with PBS for 2 times and resuspended in an equal volume of an appropriate buffer. The collected aggresomes were confirmed under a fluorescent microscope to ensure proper isolation and integrity. This protocol ensured that both the cytosolic fraction and the resuspended aggresome fraction were in equal volumes, thus maintaining consistency for subsequent analyses and allowing for accurate comparative studies in the later stages of the experiment. The samples were pre-treated with 200 μg/mL Proteinase K (Thermo Fisher, #EO0491) in 10 mM Tris-HCl buffer (pH 7.5) at 50°C for 2 hours to degrade contaminating proteins. This was followed by heat inactivation at 75°C for 15 minutes. RNA was subsequently extracted and purified from each fraction using the Bacteria RNA Extraction Kit (Vazyme, R403-01), with the substitution of RNA isolater Total RNA Extraction Reagent for RNAiso Plus reagent (Takara, #9109). The purified RNA was eluted in equal volumes of nuclease-free water. Finally, RNA concentration was measured for each fraction using a Qubit Fluorometer 3 with the Equalbit RNA HS Assay Kit, with samples appropriately diluted to fall within the 5-100 ng/μL range for accurate quantification. Samples are appropriately diluted to fall within this range, ensuring precise measurements. Throughout all processes, we maintained equal volumes to ensure consistent RNA extraction efficiency across all samples, minimizing variability caused by differences in cell numbers or liquid volumes.

### RNA/Aggresome Quantification

This process involved isolating aggresomes from an equal volume of cell lysate as used in previous measurements. Next, a specific number of aggresomes were isolated by Fluorescence-Activated Cell Sorting (FACS), and RNA was purified from these aggresomes using the Vazyme (R403-01) kit, with the substitution of RNA isolater Total RNA Extraction Reagent for RNAiso Plus reagent (Takara, #9109). The RNA/aggresome ratio was then calculated by dividing the mass of purified RNA by the number of aggresomes, providing a standardized measure of RNA content per aggresome.

### FACS analysis

The HslU protein was fused with the fluorescent protein GFP to label Aggresomes within cells. Cells were treated under various conditions according to the experimental requirement and Aggresomes were subsequently extracted. FACS analysis was performed on the Aggresome suspension using an Agilent Novocyte instrument, with the following parameters configured: FSC-H: >1000; Sample Flow Rate: High; Fluorescence Selection: FITC. Aggresome concentration and count information for the suspension was obtained.

### Aggresome Count/Cell Quantification

This process utilized fluorescence microscopy to directly count aggresomes per cell. The aggresome marker HslU was labeled with GFP, following the methods described in previous studies^9,10^. For each sample, 50 randomly selected cells were analyzed. To ensure reproducibility and statistical significance, the entire procedure was repeated in triplicate (i.e., n=3).

### Cytosolic RNA/Cell Quantification

To determine the cytosolic RNA content, we employed a subtractive method. We calculated the cytosolic RNA/Cell values by subtracting the aggresome RNA/cell from the total cellular RNA/cell.

### RNA Concentration Measurements Using a Qubit Fluorometer

The Qubit Fluorometer 3 (Q33216, Invitrogen), in conjunction with the Equalbit RNA HS Assay Kit (EQ211, Vazyme), was employed for accurate RNA concentration measurements. The working solution was prepared and aliquoted, with 10 μL of Standard #1 and 10 μL of Standard #2 each diluted into 190 μL of the working solution for machine calibration and the generation of an accurate standard curve. Given the excellent linear relationship of the assay kit within the range of 5 ng-100 ng/μL for RNA, RNA samples were diluted prior to measurement to ensure their concentrations fell within this range. For each sample, 1 μL of RNA was mixed with 199 μL of the working solution, and the mixture was loaded into the Qubit Fluorometer for automatic concentration determination based on the standard curve.

### RNA-seq

The total RNA of the samples was extracted using the Bacteria RNA Extraction Kit (R403-01, Vazyme) and subjected to mRNA selection, fragmentation, cDNA synthesis, and library preparation using the VAHTSTM Total RNA-seq (H/M/R) Library Prep Kit for Illumina® (NR603, Vazyme). The library quality was analyzed on a Bioanalyzer. High-throughput sequencing was conducted on the Genome Analyzer IIx (Illumina).

### RT-qPCR

To prepare the samples for RT-qPCR analysis, overnight cultures were 1:100 diluted into fresh LB medium and incubated in a shaker at 220 rpm for 3 hr, then treated with 2 mM NaAsO_2_ for varying lengths of time according to the experimental requirements. After treatment, the cells were harvested by centrifugation and total RNA was extracted using the Bacteria RNA Extraction Kit (Vazyme China). To synthesize cDNA, 300 ng of RNA was used with the HiScript III RT SuperMix for qPCR (+gDNA wiper) (Vazyme China), following the manufacturer’s instructions. The qPCR reactions were performed using the ChamQ Universal SYBR qPCR Master Mix (Vazyme China) and the BIO-RAD CFX Connect Real-Time PCR Detection System from Bio-Rad (USA), according to the manufacturer’s instructions. The primers for RT-qPCR analysis were listed in Supplementary Table 4. The relative amount of mRNA is determined by normalizing it to the level of the internal control gene, 16s RNA. To assess spatial transcript distribution during stress, E. coli cultures undergoing sodium arsenite stress were fractionated into aggresome-enriched and cytosolic components at two time points: 30 min and 3 hr post-stress induction. RNA was isolated from both fractions and used for qPCR measurement.

### ScreenTape assay

To conduct the ScreenTape experiment, the High Sensitivity RNA ScreenTape assay kit (Agilent, America) were utilized to achieve high accuracy in measuring RNA samples at a concentration of approximately 10 ng/µL. Prior to analysis, all RNA samples were diluted, and 1 µL of RNA buffer was thoroughly mixed with 2 µL of RNA, followed by incubation at 72 °C for 3 mins and 1 min on ice. Subsequently, individual samples were subjected to RNA length distribution analysis using the Agilent 4200 (G2991AA) TapeStation system (Agilent, America).

### Translation following stress removal assay

The highly enriched mRNA *glpK* in the aggresome was selected to perform the experiment. The pBAD vector was used as a backbone to generate a recombinant plasmid that expressing the GlpK-GFP fusion protein. Overnight bacterial cultures expressing pBAD-GlpK-GFP were 1:100 diluted into fresh LB medium and incubated at 37 °C with shaking at 220 rpm for 2 hr. 0.001% arabinose was then added to induce protein and mRNA expression of GlpK-GFP with continuous incubation for another 2 hr. Cell cultures were then treated with 2mM NaAsO_2_ and 100ng/μL Rifampicin for 30 min. The transcription of *gplK*-mRNA in cytosol was inhibited by rifampicin, quenching GlpK-GFP fluorescence with laser at 488 nm. The fluorescence recovery rate of GlpK-GFP was quantified by continuously incubating the cells at 30 °C under a Nikon N-SIM S super-resolution microscope.

### Electron microscopy

The cells were collected by centrifugation, and TEM fixative was added to the tube for fixation at 4°C. Agarose pre-embedding was performed using a 1% agarose solution, and the precipitation was suspended with forceps and wrapped in agarose before it solidified. The samples were then post-fixed by treating the agarose blocks with 1% OsO_4_ in 0.1 M PB (pH 7.4) for 2 hr at room temperature. Subsequently, the samples were rinsed in 0.1 M PB (pH 7.4) three times for 15 mins each. To dehydrate the samples, a series of ethanol solutions were used at room temperature, followed by two changes of acetone. The samples were then incubated in pure EMBed 812 for 5-8 hr at 37°C. Polymerization was achieved by placing the samples in a 65 °C oven for more than 48 hr. Next, the resin blocks were cut into 50 nm thin sections using an ultramicrotome, and the tissue sections were transferred onto 150 mesh cuprum grids with formvar film. The sections were stained with a 2% uranium acetate saturated alcohol solution, avoiding light, for 8 mins, followed by staining in a 2.6% lead citrate solution, avoiding CO_2_, for 8 mins. Finally, the cuprum grids were observed using a transmission electron microscope (Hitachi, HT7800/HT7700).

### Persister counting assay

Cultures subjected to different stress conditions were diluted to an OD600 of 0.1 in fresh LB broth. Ampicillin or kanamycin was then added at a final concentration of 10× the predetermined minimum inhibitory concentration (MIC). Cultures were then incubated with shaking (250 rpm) at 37°C for 4.5 h. Following incubation, samples were withdrawn, serially diluted in LB broth, and spot-plated onto LB agar plates. Plates were incubated overnight at 37°C. Colonies were counted manually the following day to determine viable cell counts (CFU/mL)^29^. All subsequent steps (incubation, dilution, plating, counting) were performed as described above.

### Statistical analysis

p > 0.05 was considered not significant. *P ≤ 0.05, **P ≤ 0.01, ***P ≤ 0.001, ****P ≤ 0.0001 by two-tailed Student’s t test, one-way ANOVA, or Log-rank (Mantel-Cox) test. Statistical analysis was performed in GraphPad Prism or Excel.

### RNA Fluorescence in situ hybridization (FISH)

The probes used in fluorescence in situ hybridization (FISH) were designed using the Probe Designer algorithm developed by Arjun Raj (van Oudenaarden Lab, Massachusetts Institute of Technology). Cells after different treatment were fixed with 4% formaldehyde and 2.5% glutaraldehyde for 30min at RT, washed twice with wash buffer (20% formamide in 2xSSC) then permeabilized with 70% ethanol overnight at 4℃. Cells were then washed twice with wash buffer and incubated with probe set at a final concentration of 250 nM in hybridization buffer (10% dextran sulfate, 35% deionized formamide, 0.001% of *E.coli* tRNA, 2mM vanadyl ribonucleoside complex and 1 mg/mL BSA in 2xSSC buffer) at 30℃ overnight protecting from light. Cells were washed with wash buffer 3 times to avoid the background and performed imaging.

### Slimfield imaging of Pepper RNA: HBC530 complex

Slimfield^18^ is an advanced light microscopy method which enables single molecule fluorescence detection and tracking to a lateral precision of approximately 40 nm^30^ imaging over millisecond timescales, both *in vitro* and *in vivo*. The illumination field is generated by underfilling the back aperture of a high numerical aperture (NA) objective (1.49 in this case) with collimated laser light resulting in a delimited excitation volume large enough to encompass a number of whole bacterial cells at the sample plane whose excitation intensity is greater than that of traditional epifluorescence microscopy by over three orders of magnitude. The excitation of fluorescent proteins both in single- and dual-colour bacterial cell strains enables a typical exposure time of 1-5 ms per frame^31–33^, down to a few hundred microseconds when using bright organic dyes^34, 35^. By using bespoke single-particle tracking software^36, 37^ and utilising step-wise photobleaching of dyes^38, 39^ and edge-preserving filtration^40^, the apparent diffusion coefficient and the stoichiometry of tracked molecules and molecular assemblies can be accurately determined, rendering distributions via kernel density estimation^41^.

For *in vitro* imaging or surface-immobilized mRNA, Polyethelene glycol (PEG) passivated slides were prepared similarly to Paul and Myong^42^ except that MeO-PEG-NHS and Biotin-PEG-NHS (Iris biotech PEG1165 and PEG1057 respectively) were used for passivation. Flow cells were made by attaching a PEG passivated coverslip to a PEG passivated slide via two pieces of double-sided tape with a 5 mm gap between them. The flow cell was first incubated for 5 mins at room temperature with 200 μg/mL neutravidin (Thermofisher Scientific, 31000) in Phosphate Buffered Saline (PBS) followed by a 200 μL wash with PBS to remove excess neutravidin. 100 μL of 30 pM Pepper RNA in PBS was then introduced into the to the flow cell and incubated for 10 mins at room temperature to allow binding of the 5_ˈ_ biotin on the RNA molecule to the neutravidin bound to the flow cell surface, excess PEPPER RNA was washed off with a 200 μL PBS wash. Finally, 200 μL of imaging buffer (40 mM HEPES, pH 7.4, 100 mM KCl, 5mM MgCl_2_, 1x HBC530) was washed into the flow cell before the open channel ends were sealed. A bespoke single-molecule microscope^31^ was utilised in order to image the surface immobilised Pepper RNA:HBC530 complex. Excitation by an Obis LS 50 mW 488 nm wavelength laser (attenuated to 20 mW) was reduced to 10 μm at full width half maxima in the sample plane to produce a narrow field of illumination^43^, with a mean excitation intensity of 0.25 mW/µm^2^. Images were magnified to 53nm/pixel and captured using a prime 95B sCMOS camera with a 5 ms exposure time. Image analysis was carried out on 60 fields of view from three separate slide preparations using previously described bespoke MATLAB scripts^34^.

For *in vivo* imaging, cells were grown overnight in LB at 37 °C with shaking at 180 rpm. The overnight cultures were diluted 1:100 in fresh LB and grown at 37 °C for 2 hr with shaking at 180 rpm. Arabinose was then added at a final concentration of 0.001% and the cultures were incubated for another 4 hr at 22 °C with shaking at 220 rpm. The cultures were then centrifuged at 4000g for 5 min to collect the cells. The collected cells were resuspended in imaging buffer with HBC ligand (1:100 dilution) and 0.001% arabinose and incubated at room temperature for 5-15 min. The cells were then spotted on agarose gel pads containing M9 medium with HBC ligand (1:100), 25 mM HEPES (pH 7.4), 5 mM MgSO_4_, 20mM arsenite and 0.01% arabinose. The cells were then imaged on a Slimfield microscope set up using the same imaging settings as for *in vitro* imaging and analysed using the same tracking software.

### Modeling

Aggresome formation was interpreted as LLPS involving a ternary mixture of protein, mRNA and aqueous solvent, driven by net gain in attractive enthalpy mediated from increased protein and RNA interactions which outweigh the loss in entropy due to a more ordered phase-separated state in the aggresome compared to solvated protein and RNA outside the aggresome, using a Flory-Huggins (FH) theory for ternary mixtures. This model should be used with care, as it invokes a mean-field approximation that negates the explicit molecular structure of the constituents – FH theory breaks down near the critical point, and it ignores impact of local correlations (such as chain-connectivity and self-avoidance^44^, which may lead to overestimates of the concentration in the dense phase^44, 45^). Nevertheless, FH theory is invaluable in the prediction of liquid-liquid phase separation in binary and multi-component mixtures; it and qualitatively describes how the propensity of this phenomenon is affected by the molecular weight and the (effective) strength of molecular interactions. The enthalpic component embodied in the χ interaction parameter depicts overall enthalpy as the sum of interactions involving the water solvent, protein, and RNA while the entropy component characterizes biomolecular mixing and translational effects. In the limit of low RNA concentrations, as suggested from our earlier measurements indicating a protein-rich aggresome environment^9^, χ is dominated by protein-solvent interactions. We assume that in the low RNA concentration limit RNA can partition into aggresomes but has negligible influence on phase separation. The enthalpic component of RNA-protein and RNA-water interactions per nucleotide base are assumed to be non-specific. For modeling RNA degradation effects, RNA diffusion outside the aggresome was modeled using the Stokes-Einstein relation assuming a length-dependent radius of gyration as predicted by a freely-jointed chain. For simplicity, RNA degradation was modeled with a fixed probability density at each nucleotide base. The probability of RNA escape from an aggresome once incorporated was modeled as a Boltzmann factor incorporating a length-dependent enthalpic factor. Full details in Supplementary Information.

### Protein surface charge analysis

For the surface charge analysis of proteins mentioned in this study, the structure of relative proteins was downloaded from Uniprot in AlphaFold format and imported into PyMOL (http://www.pymol.org), a user-sponsored molecular visualization system on an open-source foundation, to calculate surface charge using the vacuum electrostatics generation function. For the mutants, surface charges were analyzed based on wild-type proteins. Relative amino acids were mutated using the mutagenesis function of PyMoL and recalculated to obtain the surface charges of mutants.

### Bioinformatics analysis

For the RNA-Seq analysis, the raw data quality of the prepared libraries was assessed using FastQC (http://www.bioinformatics.babraham.ac.uk/projects/fastqc/). The reads were mapped to the MG1655 k12 genome (Ensembl Bacteria, Taxonomy ID: 511145) using the BWA aligner software (version 0.7.17-r1188, https://github.com/lh3/bwa.git). Then the sam files was converted to bam files using samtool (Version 1.9, https://samtools.sourceforge.net/). The bam files were counted with featureCounts (Version 2.0.1, https://github.com/topics/featurecounts) to generate expression results. Differential expression analysis was performed using DESeq2^46^. For Gene Ontology (GO) analysis and Kyoto Encyclopedia of Genes and Genomes (KEGG) pathway analysis, gene id was converted using the DAVID web resource (http://david.abcc.ncifcrf.gov/). R package clusterProfiler R package^47^ was used to identify the pathways in which these up- and down-regulate genes are enriched.

For distribution of reads mapping to the 3’ and 5’ fragments of operons in the cytosol and aggresomes, each transcript is bisected into 3’ and 5’ fragments. The reads from next-generation RNA sequencing are mapped to these 3’ and 5’ fragments (Ratio (3’ fragment) = 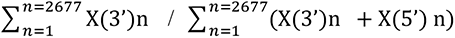, Ratio (5’ fragment) = 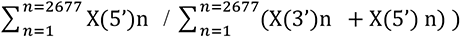, illustrating the relative abundance of each fragment in the respective cellular compartments. X(3’)n and X(5’)n represent the reads obtained from next-generation RNA sequencing, which are aligned to the 3’ and 5’ fragments of each transcript, respectively. The information regarding transcripts (possibly multiple transcripts in one operon) has been sourced from the EcoCyc *E. coli* database available at https://ecocyc.org/.

For the mass spectrometry analysis, aggresome and cytosol proteins were analyzed using mass spectrometry (MS), and the resulting data were first normalized by the total number of proteins per sample. Then, differential analysis was performed using DESeq2 to find the differentially expressed proteins. Gene Ontology (GO) analysis was performed using clusterProfiler R package for the enriched proteins of the samples. Protein-protein interaction network functional enrichment was then analyzed using STRING^48^ (https://cn.string-db.org/) for proteins enriched in aggresome or cytosol with an average expression abundance greater than 50. In addition to this, the GeneMANIA prediction server (biological network integration for gene prioritization and predicting gene function. http://pages.genemania.org/) was used to analyze the proteins. The interaction data were exported and later plotted using Cytoscape (An open-source platform for complex analysis and visualization. https://cytoscape.org/).

### RNase enzyme activity measurement

We purified RNB-WT RNB-ME and RNB-MO respectively and used the RNase Viability Assay Kit (Fluorescent Labeling) (Yeasen, 41309ES96) to measure the RNase enzyme activity. This method utilizes a unique RNA substrate, with one end labeled with a Fluorophore (Fluor) and the other end labeled with a quenching group. In the absence of RNase, the physical proximity of the quenching groups suppresses the fluorescence of the fluorophore to a low level. When RNase is present, the RNA substrate is hydrolysed, causing the spatial separation of the fluorophore and the quencher, resulting in bright fluorescence emissions. After adding the protein sample to the reaction solution, explicitly following manufacturer’s protocol, we firstly record the initial fluorescence value RFU0, then incubate at 37℃ for 1 hour, and record the fluorescence value RFU60. The ratio of RFU60/RFU0 is a metric for whether the sample has RNase activity or not: an RFU60/RFU0>2 indicates the presence of RNase activity in the sample. The results are shown in Extended Figure 9d. Negative ctrl: Nuclease-free H2O. Positive control: RNaseA.

## DATA AVAILABILITY STATEMENT

RNA-seq data are publicly available via GEO under accession GSE293685. Mass spectrometry data are deposited in iProX under accession PXD065481. Additional datasets are accessible via Zenodo at doi.org/10.5281/zenodo.15738775 and doi.org/10.5281/zenodo.12758836.

## CODE AVAILABILITY STATEMENT

All bioinformatic analysis code is publicly hosted on GitHub: RNA-seq pipeline (github.com/123456yxd/Code-of-RNA-seq, doi.org/10.5281/zenodo.15803504), biophysics analysis (github.com/elifesciences-publications/york-biophysics, doi.org/10.5281/zenodo.15805285), and aggresome modeling (github.com/CharleySchaefer/AggresomeIPBM, doi.org/10.5281/zenodo.15806185).

## ACKNOWLEDGMENTS

This work was supported by the grants from the Major Project of Guangzhou National Laboratory (GZNL2024A01023), the Fundamental Research Funds for the Central Universities (2042022dx0003), Natural Science Foundation of Wuhan (2024040701010031), National Natural Science Foundation of China (31970089, T2125002, 82241230, 82341007), National Key R&D Program of China (2021YFC2701602, 2022YFC2504602), the Engineering and Physical Science Research Council (EP/W024063/1, EP/Y000501/1) and Biotechnology and Biological Sciences Research Council (BB/W000555/), the Beijing Natural Science Foundation (Z220014), the New Cornerstone Science Foundation through the XPLORER PRIZE.. We also thank all the staff in the Core Facilities of Medical Research Institute at Wuhan University and the Core Facilities at School of Life Sciences at Peking University for their technical support.

## AUTHOR CONTRIBUTION STATEMENT

Conceptualization: Y.P., F.B., M.L.; Methodology: L.P., Y.X., X.Y., C.S., A.S., J.H., H.L.; Investigation: L.P., Y.X., X.D., C.S., A.S., J.H.; Bioinformatics analysis: X.Y., W.Z.; Supervision: Y.P., M.L.; Writing – original draft: Y.P., M.L.; Writing – review & editing: Y.P., M.L., F.B.

---

**Extended Data Fig. 1.**
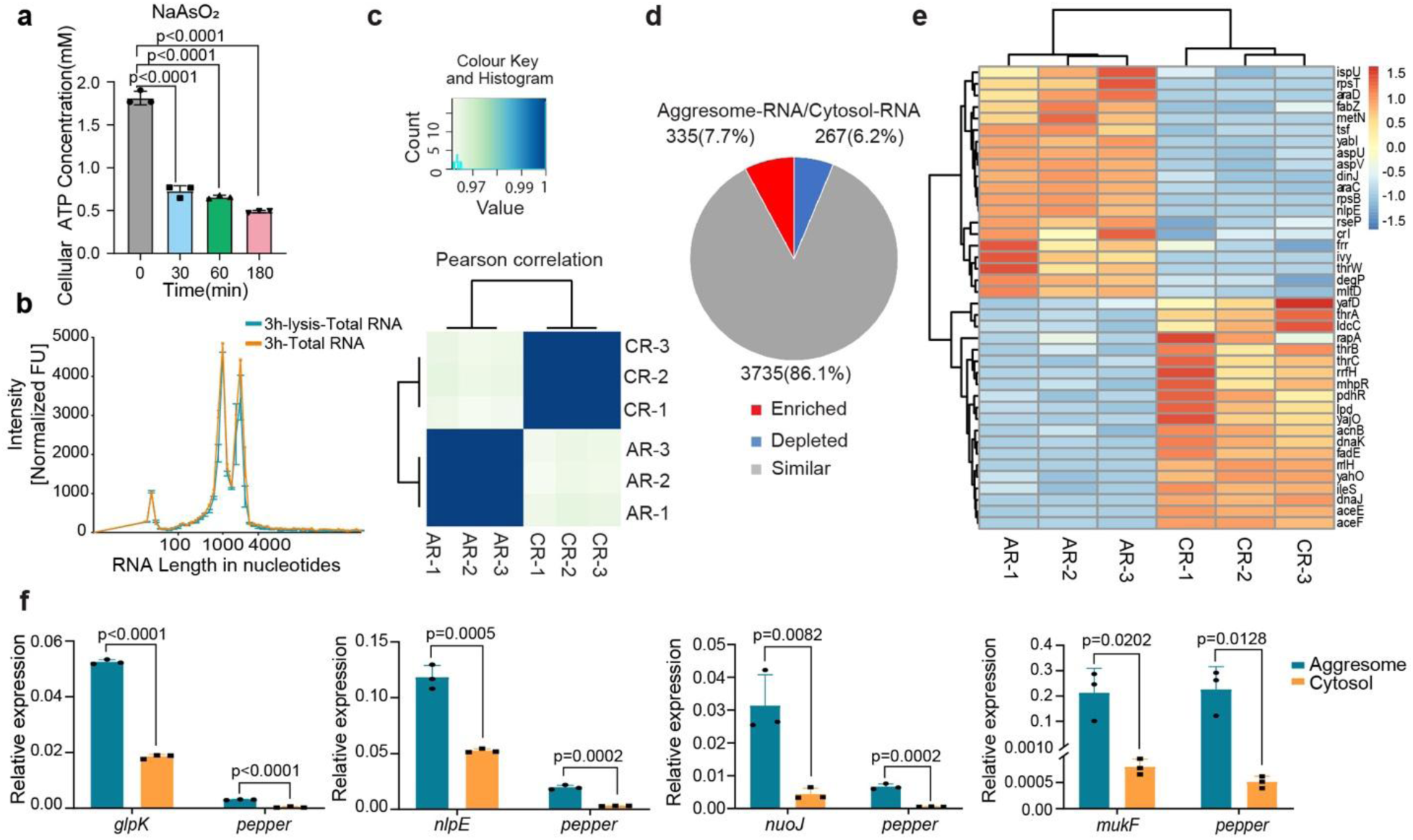
| Aggresome formation enriches mRNA. **a**. Cellular ATP concentration after arsenite (2mM) treatment for various time durations (n=3 independent biological replicates, mean ± SE). **b.** RNA length distributions determined by ScreenTape analysis: *3h-total RNA from whole cells*: Standard extraction from exponential-phase cells. *Total RNA from cell lysate*: Lysate prepared via adjusted protocol prior to standard extraction (n=3 independent biological replicates, mean ± SE). **c**. Pairwise correlation coefficients between Aggresome-RNA library duplicates and Cytosol-RNA library duplicates, indicating that the aggresome transcriptome is distinct from that of the cytosol (Pearson correlation coefficient, R_2_ < 0.001). **d**. Pie chart depicting gene number and the relative contribution of each class of RNA (Aggresome enriched, Aggresome depleted, or neither) to the cytosol transcriptome. **e**. Heatmap showing relative transcript abundance of Aggresome-RNA and Cytosol-RNA. Scale beside the heatmap indicates log2-normalized transcript abundance relative to the mean expression level (n=3 independent biological replicates). **f**. mRNA expression level of the genes measured by quantitative RT-PCR (n=3 independent biological replicates, mean ± SE). Two-sided unpaired Student’s t-test used in comparison; error bars indicate SE.

**Extended Data Fig. 2.**
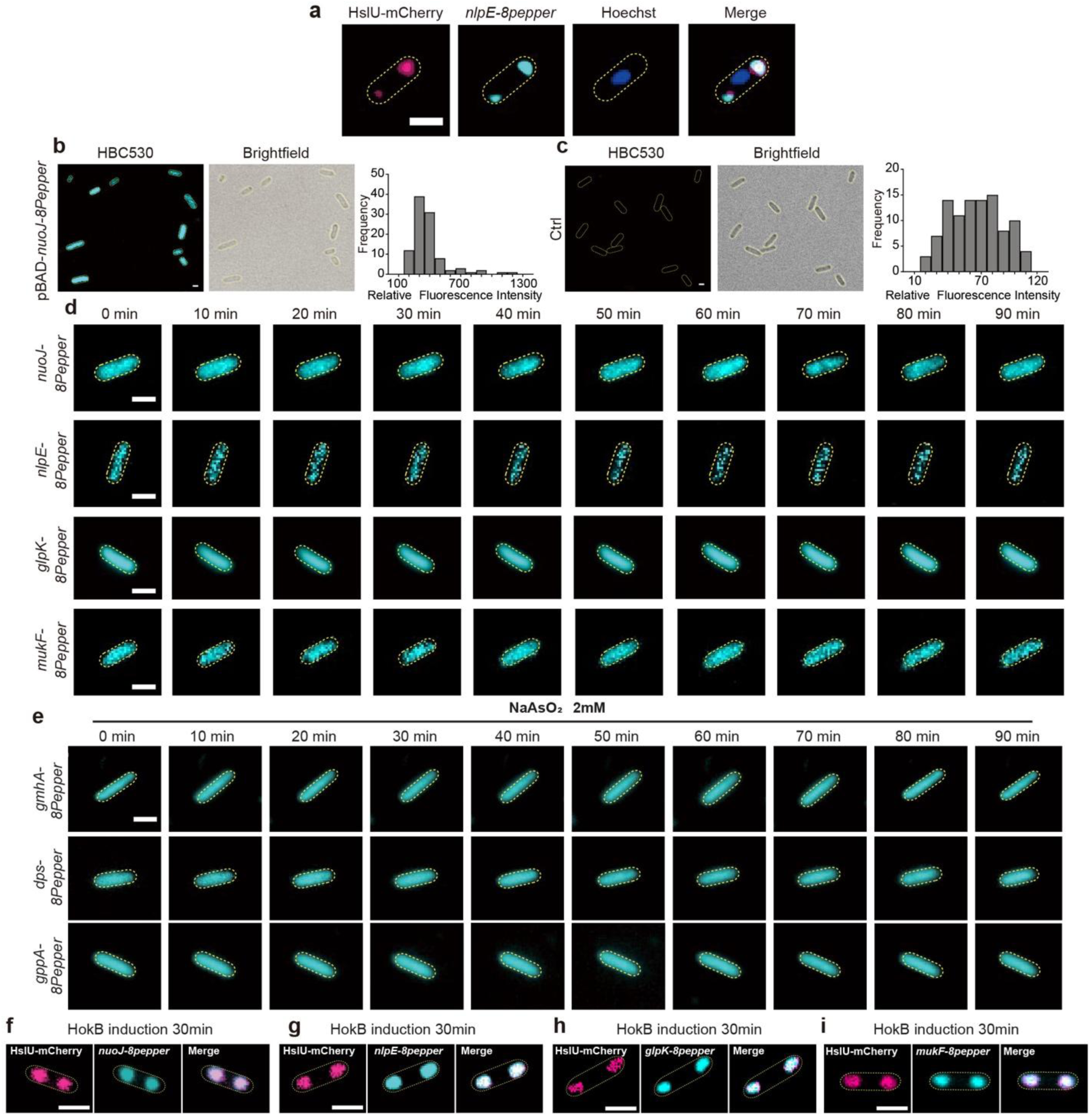
| mRNA localization to bacterial aggresomes. **a.** Epifluorescence image of aggresome (induced by 2 mM arsenite, 30 min) showing colocalization of nlpE mRNA (nlpE-8pepper/HBC530), protein (HslU-EGFP), and DNA (Hoechst). **b,c.** Control images: **(b)** Live E. coli expressing Pepper aptamer stained with HBC530. **(c)** Wild-type E. coli stained with HBC530 (1 μM). **d.** Distribution of glpK-8pepper mRNA in unstressed cells. **e.** Distribution of gmhA/dps/gppA-8pepper mRNAs in cells under arsenite treatment (2 mM, 30 min). **f-i.** SIM imaging showing aggresomal partitioning of nuoJ-8pepper (**f**), nlpE-8pepper (**g**), glpK-8pepper (**h**), and mukF- 8pepper (**i**) mRNAs under HokB induction (30 min). All mRNAs labeled via 8pepper/HBC530 in imaging buffer. Scale bars: 1 μm.

**Extended Data Fig. 3.**
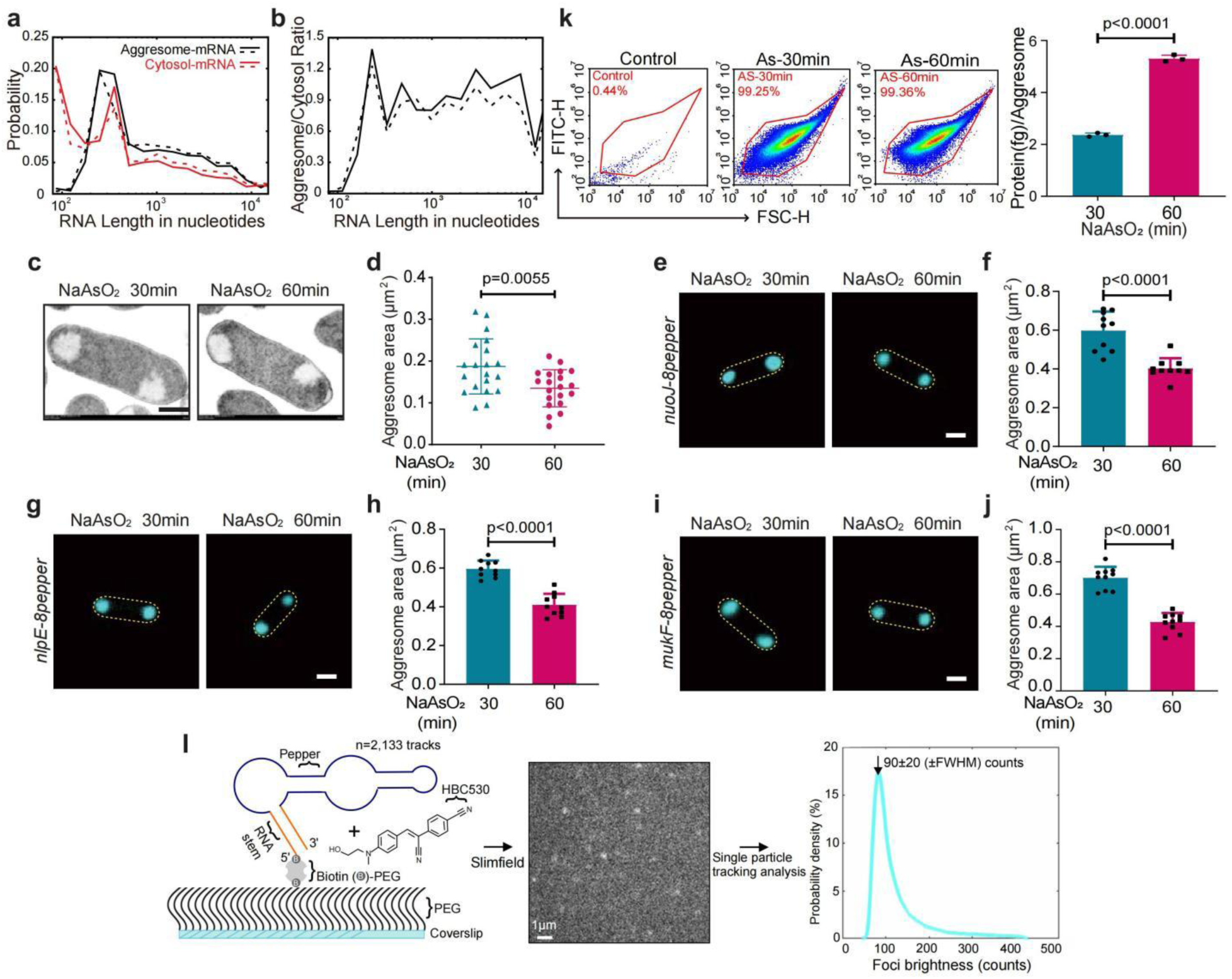
| RNA characterization of aggresomes. **a.** Distribution of RNA lengths in aggresomes versus cytoplasm, analyzed using transcriptome sequencing data aligned to operonic mRNA references. **b.** Aggresome-to- cytoplasm RNA ratio as a function of transcript length (nucleotides), derived from transcriptome sequencing data referenced against operonic mRNAs. **c, d.** Aggresome morphology by transmission electron microscopy (TEM): **c.** Representative TEM images post-arsenite treatment. Scale bar: 500 nm. **d.** Aggresome area quantification from TEM data (n = 20 cells per condition, mean ± SE). **e–j.** Analysis of aggresome compaction via SIM: **e, g, i.** SIM images of cells expressing *nuoJ*-8pepper (**e**), *nlpE*-8pepper (**g**), or *mukF*-8pepper (**i**) after indicated arsenite (NaAsO_2_) treatment durations. HBC530 dye was used for RNA visualization. Scale bars: 1 µm. **f, h, j.** Aggresome area quantification from **e**, **g**, and **i**, respectively (n = 10 cells per condition). **k.** Protein mass per aggresome after arsenite treatment: *Protein total*: Bulk aggresome protein (Qubit fluorometry). *N*_aggresome_=Aggresome count (FACS). Protein per aggresome = *Protein total* / *N*_aggresom_ (n = 3 independent biological replicates, mean ± SE). **l.** Workflow for *in vitro* single-molecule mRNA detection: *Left*: Pepper RNA (stem: orange; aptamer: blue) immobilized on a passivated coverslip and incubated with HBC530 dye. *Middle*: Slimfield microscopy localizes dye-bound complexes as diffraction-limited foci (ms timescale). *Right*: Custom single-particle tracking software determines centroid positions (∼40 nm precision) and quantifies focus brightness (modal intensity: ∼90 counts; background-subtracted). Two-sided unpaired Student’s t-test used in comparison; error bars indicate SE.

**Extended Data Fig. 4.**
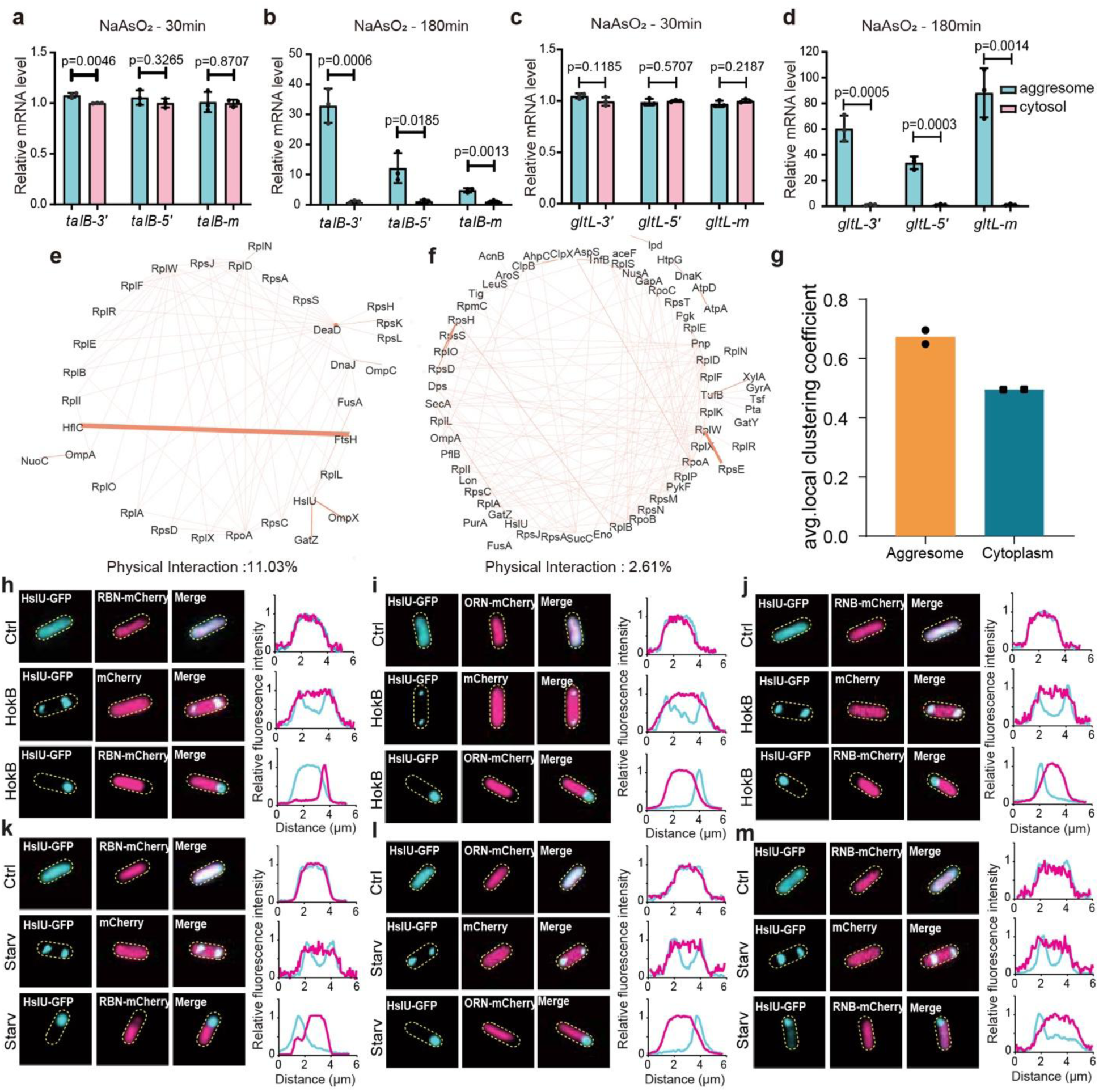
| Protein interaction networks and ribonuclease localization under stress conditions. a-d. Relative RNA levels of representative transcripts (*talB*, *gltL*) in aggresomes versus cytosol after 30 min or 180 min of 2 mM arsenite treatment (n = 3 independent biological replicates). **e, f.** Protein-protein interaction (PPI) networks: **e.** Aggresome-associated proteins. **f.** Cytoplasmic proteins. **g.** Comparison of average local clustering coefficients between aggresome and cytoplasmic protein networks (n=2 independent biological replicates). **h-m.** Subcellular distribution of ribonucleases under stress: **h-j.** Localization of RBN-mCherry (**h**), ORN-mCherry (**i**), and RNB-mCherry (**j**) during HokB toxin induction. **k-m.** Localization of RBN-mCherry (**k**), ORN-mCherry (**l**), and RNB-mCherry (**m**) during starvation. *For all panels:* Aggresomes marked by HslU-GFP. Right: Fluorescence intensity profiles of GFP (aggresome) and mCherry (ribonuclease) along the cellular long axis. Conditions: Ctrl (exponential phase, untreated), Ars (2 mM arsenite). Scale bar: 1 μm. Two-sided unpaired Student’s t-test used in comparison; error bars indicate SE.

**Extended Data Fig. 5.**
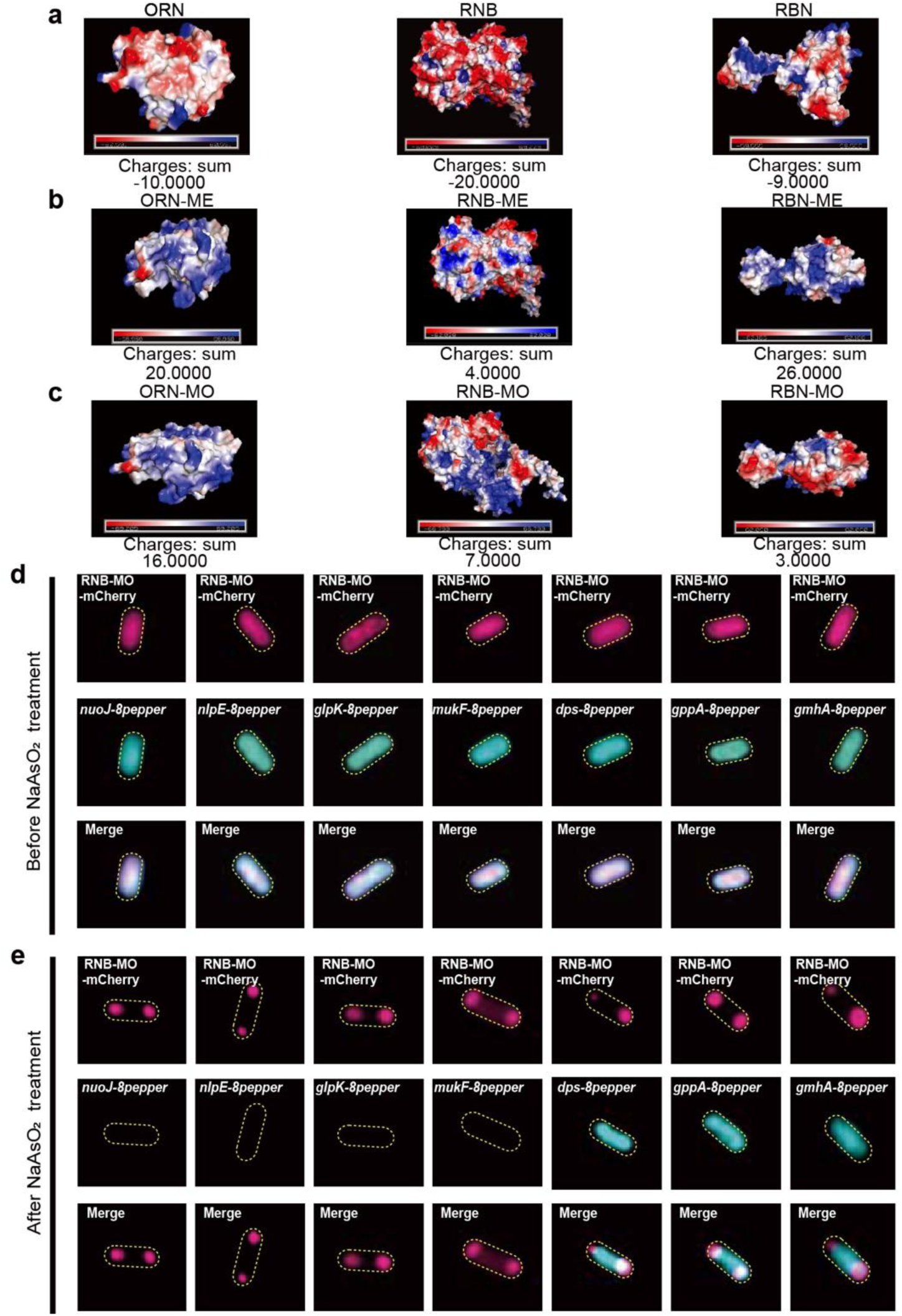
| Ribonuclease surface properties and stress-induced mRNA localizatio. a-c. Protein surface charge analysis (PyMOL): **a.** Wild-type ribonucleases (ORN, RNB, RBN). **b.** ME mutants: Alanine substitutions at all D/E residues within enzymatic centers. **c.** MO mutants: Alanine substitutions at D/E residues outside RNA-binding motifs/catalytic centers. **d-e.** Subcellular mRNA distribution (SIM): **d.** 8pepper-tagged mRNAs (*nuoJ, nlpE, glpK, mukF, gmhA, dps, gppA*) in untreated cells. **e.** Same mRNAs after 30-min 2 mM arsenite treatment. *Imaging:* HBC dye in imaging buffer. Scale bar: 1 μm

**Extended Data Fig. 6.**
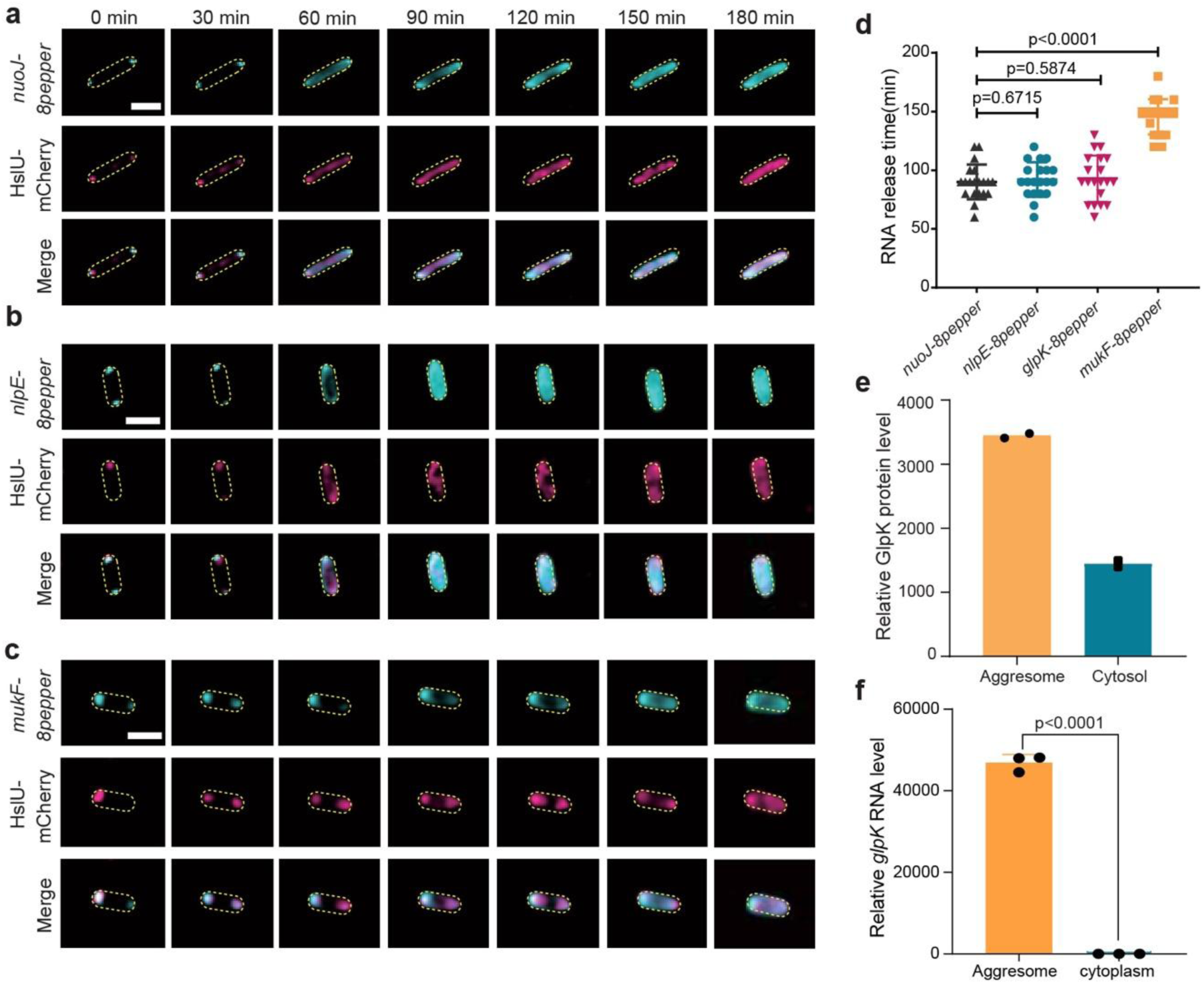
| mRNA release kinetics and compartment-specific molecular levels post-arsenite removal. a-c. Release of 8pepper-tagged *nuoJ* (**a**), *nlpE* (**b**), and *mukF* (**c**) mRNAs from aggresomes (marked by HslU-mCherry) following arsenite washout. **d.** Mean mRNA release duration for each transcript (n=20 tracked cells, mean ± SE). **e.** Relative GlpK *protein* levels in aggresomes vs. cytoplasm post-arsenite treatment (quantified by mass spectrometry, MS; n=2 independent biological replicates). **f.** Relative *glpK RNA* levels in aggresomes vs. cytoplasm post-arsenite treatment (RNA-seq; n=3 independent biological replicates, mean ± SE). Imaging for **a-c:** HBC dye in imaging buffer. Scale bar: 2 μm. Two-sided unpaired Student’s t-test used in comparison; Error bars = SE.

**Extended Data Fig. 7.**
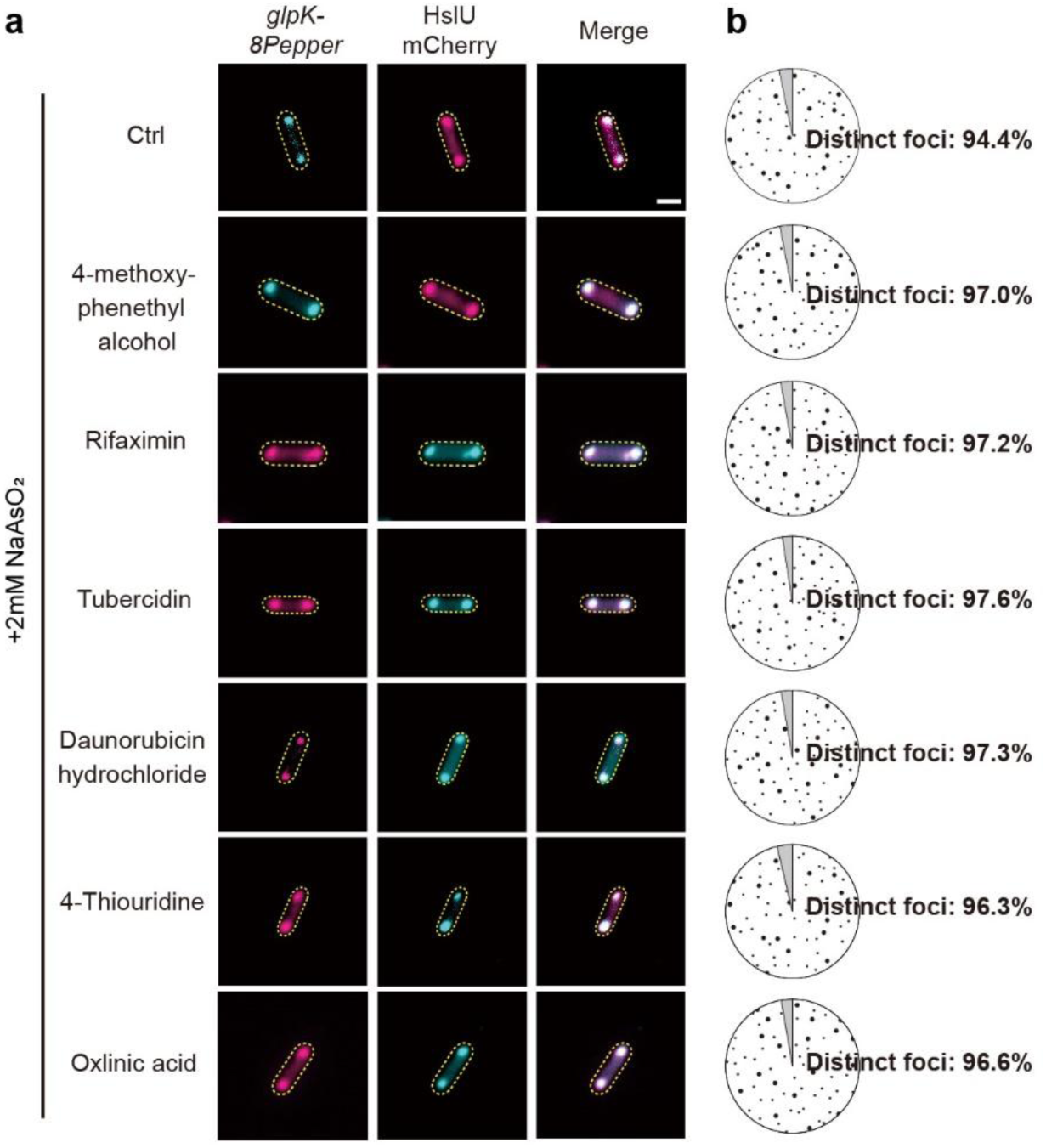
**| Small-molecule screen for inhibitors of RNA recruitment to aggresomes**. **a.** Representative fluorescence microscopy images of cells treated with different chemical combinations. Aggresomes visualized as distinct mRNA foci. **b.** Quantification of cells containing aggresomes (distinct mRNA foci) across treatment conditions. Data derived from **a** (n = 50 cells per condition; 3 independent biological replicates).

