## Supplementary Information for "Aggresomes protect mRNA under stress in *Escherichia coli*"

### I. SUPPLEMENTARY NOTE 1: INTRODUCTION

In this section, we will interpret the observation of droplet compaction and the partitioning of RNA. The key assumption is that these processes take place within a sufficiently short time for the number of RNA nucleotides to be conserved (i.e., no RNA is expressed, and no RNA diffuses out of the cell). Secondly, we will describe droplet compaction and RNA partitioning using a Flory-Huggins (FH) model for ternary mixtures. This model invokes a mean-field approximation that fails near critical points, but which allows for physically meaningful predictions away from criticality (see Ref. 1). In the following, we first present the FH model for a ternary mixture of protein, RNA, and solvent. We then discuss the regime of low RNA concentrations, where phase separation is purely driven by the incompatibility between the protein and the solvent, described by the $\chi_{\mathrm{PS}}$ interaction parameter. We show through a mass balance of a closed system (i.e., the cell), that an increased $\chi_{\mathrm{PS}}$ parameter leads to droplet compaction: the droplet size decreases and its internal protein concentration increases. We then separately show using the same ternary FH model that RNA partitioning can be explained using non-specific RNA-solvent and RNA-protein interaction parameters, with a preference of RNA to interaction with the protein rather than the solvent. The theory predicts that RNA partitions stronger with an increasing chain length, in quantitative agreement with ScreenTape experiments.

### II. SUPPLEMENTARY NOTE 2: TERNARY FLORY-HUGGINS MODEL

We interpret the liquid-liquid phase separation of protein-rich aggresomes and the attraction of RNA molecules using a ternary Flory-Huggins theory for the free energy per segment[2],

$$\begin{aligned} f/k_{\mathrm{B}}T = & \left(\frac{1}{N_{\mathrm{P}}}\phi_{\mathrm{P}}\ln\phi_{\mathrm{P}} + \phi_{\mathrm{S}}\ln\phi_{\mathrm{S}} + \phi_{\mathrm{P}}\phi_{\mathrm{S}}\chi_{\mathrm{PS}}\right) \\ & +\phi_{\mathrm{R}}\left(\frac{1}{N_{\mathrm{R}}}\ln\phi_{\mathrm{R}} + \chi_{\mathrm{RS}}\phi_{\mathrm{S}} + \chi_{\mathrm{RP}}\phi_{\mathrm{P}}\right), \end{aligned} \tag{1}$$

with $k_{\mathrm{B}}T$ the thermal energy, $\phi_{\mathrm{S}}$, $\phi_{\mathrm{P}}$, $\phi_{\mathrm{R}}$ the volume fractions of solvent, protein, and RNA, respectively, and with

$$\chi_{AB} = \frac{z}{k_{\mathrm{B}}T}\left(\varepsilon_{AB} - \frac{1}{2}(\varepsilon_{AA} + \varepsilon_{BB})\right) \tag{2}$$

the Flory-Huggins interaction parameter between components $A$ and $B$ for $A, B = \mathrm{R}, \mathrm{P}, \mathrm{S}$, with $z$ the coordination number and the $\varepsilon$s the interaction energies. Further, $N_{\mathrm{R}}$ and $N_{\mathrm{P}}$ are dimensionless measures for the molecular weights of RNA and the protein, respectively, and they control the (translational) mixing entropy of the molecules. We have arranged the terms into a part that describes the translational entropy of the protein and the solvent and the protein-solvent interactions, and a part that describes the translational entropy of the RNA molecule and the RNA-protein and RNA-solvent interactions.

### III. SUPPLEMENTARY NOTE 3: AGGRESOME COMPACTION

In the limit of low RNA concentrations, the terms within the right set of brackets can be ignored, and the free energy reduces to the binary expression $f_{\mathrm{SP}}/k_{\mathrm{B}}T = \phi_{\mathrm{S}}\ln\phi_{\mathrm{S}} + \frac{1}{N_{\mathrm{P}}}\phi_{\mathrm{P}}\ln\phi_{\mathrm{P}} + \phi_{\mathrm{S}}\phi_{\mathrm{P}}\chi_{SP}$. This expression predicts liquid-liquid phase separation between the protein and the solvent if the interaction parameter is larger than the critical value

$$\chi_{\mathrm{SP}} > \chi_{\mathrm{SP,cr}} = \frac{1}{2}\left(1 + \frac{1}{\sqrt{N_{\mathrm{P}}}}\right)^2, \tag{3}$$

and if the concentration is within the range bound by the binodal concentrations, which are found using the a common tangent construction [2]; see blue curve (calculated for $N_{\mathrm{P}} = 50$) in the left panel of Figure 1. The binodal describes the protein concentration in the cytosol, $\phi_{\mathrm{P,C}}$ in coexistence with the protein concentration in the aggresome, $\phi_{\mathrm{P,A}}$, and is related to the overall concentration $\phi_{\mathrm{P}}$ through the mass balance

$$\phi_{\mathrm{P}}(V_{\mathrm{C}} + V_{\mathrm{A}}) = \phi_{\mathrm{P,A}}V_{\mathrm{A}} + \phi_{\mathrm{P,C}}V_{\mathrm{C}}, \tag{4}$$

in which $V_{\mathrm{A}}$ is the total volume of all aggresome droplets and $V_{\mathrm{C}}$ is the effective volume of the cytosol (i.e., it includes any volume within the cell that is not within the aggresome). As a consequence of this mass balance with constant

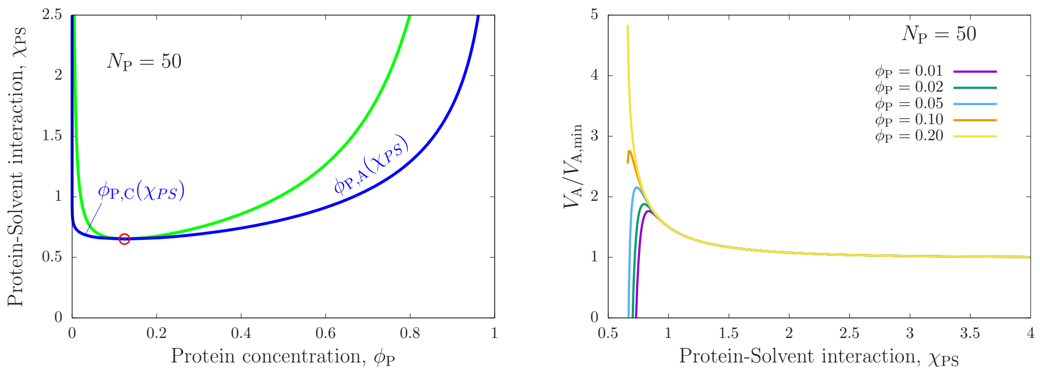


FIG. 1. We rationalise liquid-liquid phase separation of droplets and their compaction in terms of a binary Flory-Huggins phase diagram for the solvent and the protein, and we use a protein size $N_P = 50$ in our qualitative discussions. Left: Phase diagram of protein and water demixing. The red circle is the critical point, and the green and blue curves represent the spinodal and binodal, respectively. The binodal gives the concentrations in the cytosol, $\phi_{P,C}$ and in the aggresome, $\phi_{P,A}$. The increase of $\phi_{P,A}$ with an increasingly protein-solvent interaction parameter, $\chi_{PS}$, combined with the mass balance constraint in in Eq. 4 causes compaction of the aggresome size displayed in the right panel. Right: The ratio between the aggresome size in a swollen state ($\phi_{P,A} \ll 1$) compared to the fully compact state ($\phi_{P,A}$) decreases with an increasingly repulsive protein-solvent interaction parameter. For protein concentrations lower than the critical value (approximately 0.134, see Left panel), first an increase of the droplet size occurs. At higher concentrations the compaction behaviour is monotonous, and qualitatively describes our experimental observations.

$V_A + V_C$, the volume of the aggresome depends on the interaction parameter through the equality

$$\frac{V_A(\chi_{SP}, \phi_P, N_P)}{V_A + V_C} = \frac{\phi_P - \phi_{P,C}(\chi_{PS})}{\phi_{P,A}(\chi_{PS}) - \phi_{P,C}(\chi_{PS})} \tag{5}$$

For high values of the interaction parameter, the protein concentration inside the aggresome tends to the maximum concentration ($\phi_{P,A} = 1$), and in turn the volume of the aggresome shrinks towards the most compact size $V_{A,\min} \equiv \phi_P(V_A + V_C)$. This dependence of the aggresome size on the interaction parameter is displayed in the right panel of Figure 1.

For low overall protein concentrations, Figure 1 shows a non-monotonous dependence. For low interaction parameters, both the size and internal concentration of the aggresome increase due to the recruitment of more proteins from the cytosol. For a further increase of the interaction parameter, the proteins in the cytosol deplete and the further increase of the protein concentration inside the aggresome is governed through droplet compaction. These two regimes exist for overall concentrations of protein lower than the critical value

$$\phi_{R,\mathrm{cr}} = \frac{1}{1 + \sqrt{N_P}}, \tag{6}$$

while for higher concentrations the initial growth stage does not occur. We thus interpret the experimental observation of droplet compaction as a consequence of an increase in the protein-solvent interaction parameter due to induction of cell stress, where the monotonous shrinkage is cause by either a a protein concentration compare to the critical concentration, or due to the protein-solvent interaction parameter being well above the critical value in Eq. 3.

### IV. SUPPLEMENTARY NOTE 4: LENGTH-DEPENDENT PARTITIONING OF RNA

We interpret RNA partitioning as a subdominant effect in response the liquid-liquid phase separation. That is, the association of RNA to the droplets has a negligible influence on phase separation. The free energy of RNA partitioning may be extracted from the full ternary free energy above,

$$f_R/k_BT = \phi_R \left( \frac{1}{N_R} \ln \phi_R + \frac{H}{k_BT} \right), \tag{7}$$

where we have defined the effective enthalpy of association to the aggresome per residue as

$$H \equiv k_BT \left( \phi_S \chi_{SR} + \phi_P \chi_{PR} \right). \tag{8}$$

This binding enthalpy depends on the volume fraction of proteins in the aggresome (note that the volume fraction of solvent is $\phi_{\mathrm{S}} \approx (1-\phi_{\mathrm{P}})$), and is thus expected to correlate with aggresome compaction.

To describe how RNA partitions between the cytosol with low protein concentration $\phi_{\mathrm{P,C}}$, and in the aggresome with high concentration $\phi_{\mathrm{P,A}}$; both given by the binodal branch in the phase diagram of Figure 1, we parameterize the steady-state condition by formulating the total free energy associated to the RNA as

$$F = V_{\mathrm{C}} f_{\mathrm{R}}(\phi_{\mathrm{R,C}}) + V_{\mathrm{A}} f_{\mathrm{R}}(\phi_{\mathrm{R,A}}), \tag{9}$$

with $V_{\mathrm{C}}$ the effective volume of the cytosol and $V_{\mathrm{A}}$ the volume of the aggresome. The RNA concentration in the cytosol, $\phi_{\mathrm{R,C}}$ and in the aggresome $\phi_{\mathrm{R,A}}$ are constrained by the mass balance $\phi_{\mathrm{R,C}}V_{\mathrm{C}} + \phi_{\mathrm{R,A}}V_{\mathrm{A}} = \phi_{\mathrm{R,tot}}(V_{\mathrm{C}} + V_{\mathrm{A}})$, where $\phi_{\mathrm{R,tot}}$ is the overall concentration of RNA. Thus, to obtain the concentrations of RNA inside and outside the aggresome, we minimise $F$ with respect to $\phi_{\mathrm{R,C}}$, with $\phi_{\mathrm{R,A}}$ dependent on $\phi_{\mathrm{R,C}}$ according to the mass balance.

$$0 = \frac{\partial F}{\partial \phi_{\mathrm{R,C}}} = V_{\mathrm{C}}\left(\frac{\partial f_{\mathrm{R}}}{\partial \phi_{\mathrm{R,C}}} - \frac{\partial f_{\mathrm{R}}}{\partial \phi_{R,A}}\right) = \frac{H_{\mathrm{C}} - H_{\mathrm{A}}}{k_{\mathrm{B}}T} + \frac{1}{N_{\mathrm{R}}}\ln \phi_{\mathrm{R,A}} - \frac{1}{N_{\mathrm{R}}}\ln \phi_{\mathrm{R,C}}, \tag{10}$$

which gives

$$\frac{\phi_{\mathrm{A}}}{\phi_{\mathrm{C}}} = \exp\left(-\frac{N_{\mathrm{R}}\Delta H}{k_{\mathrm{B}}T}\right) \tag{11}$$

as the ratio between the RNA concentrations outside and inside the aggresome. The difference in enthalpies is

$$\Delta H = \Delta\phi_{\mathrm{P}} k_{\mathrm{B}} T(\chi_{\mathrm{RP}} - \chi_{\mathrm{RS}}), \tag{12}$$

with $\Delta\phi_{\mathrm{P}} = \phi_{\mathrm{P,A}} - \phi_{\mathrm{P,C}}$ the concentration difference of the proteins inside and outside the aggresome.

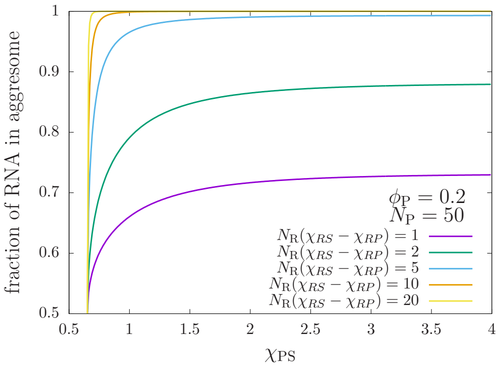


FIG. 2. Fraction of RNA inside the aggresome against the interaction strength between the protein and solvent, plotted for various length of the RNA chain $N_{\mathrm{P}}$ (scaled using the difference in interaction parameters $\chi_{\mathrm{RS}}$ and $\chi_{\mathrm{RP}}$). The relatively long RNA chains are efficiently partitioned at modest values of the protein-solvent interaction parameter $\chi_{\mathrm{PS}}$, while the recruitment of shorter RNA chains requires higher interaction strengths. This is due to an increased number of RNA-protein interactions, because of the increased protein concentration at high $\chi_{\mathrm{PS}}$ values, see phase diagram in Fig. 1.

A small Flory-Huggins $\chi$ parameter implies a good compatibility of two components. Hence, if $\chi_{\mathrm{RP}}$ is smaller than $\chi_{\mathrm{RS}}$, the RNA is more compatible with the protein than with the solvent, and, thus, results in a negative enthalpy of RNA association to the aggresome, $\Delta H$, as expected. Indeed, for negative $\Delta H$ the RNA concentration inside the aggresome becomes high compared to the outside concentration. This partitioning effect becomes stronger with an increasing number of residues, which we indeed find in the experimental data shown in Fig 2 of the main text. The partitioning also becomes stronger for more negative values of $\Delta H$, which may happen if the concentration of protein increases inside the aggresome, as described by Eq. 12. We indeed measured this effect in response to cell stress induction using a range of stress factors, where aggresome droplets shrink and the inside protein concentration increased, and in response recruited a higher RNA content.

*Application to ScreenTape data.* – As a proxy for the concentration of the RNA molecules, dependent on the residues $N_{\mathrm{R}}$, inside and outside the aggresome we have used the screen tape intensities $I_{\mathrm{AR}}(N_{\mathrm{R}})$ and $I_{\mathrm{CR}}(N_{\mathrm{R}})$, respectively. We have used only the data with intensities well above a background intensity that we quantified used in the control intensity, $I_{\text{control}}(N_{\text{R}})$ as follows: We used the normalised error of the control intensity averaged for all datapoints as a tolerance measure,

$$\text{tolerance} = \left\langle \frac{\sigma_{I_{\text{control}}}}{I_{\text{control}}} \right\rangle, \tag{13}$$

and averaged the $I_{\text{control}}$ intensities from $N_{\text{R}} = 6654.7$ and up to get the baseline value. This baseline value was chosen such that the normalised error around the baseline matched the tolerance value (for smaller $N_{\text{R}}$ the normalised error increased due to the inclusion of intensities above the mean of the baseline). The baseline and the error on the baseline was then used to exclude any data with $I_{\text{AR}} - \sigma_{I_{\text{AR}}} < I_{\text{control}} + \sigma_{I_{\text{control}}}$ and $I_{\text{CR}} - \sigma_{I_{\text{CR}}} < I_{\text{control}} + \sigma_{I_{\text{control}}}$. The data that remained after filtering was used to calculate the ratio $I_{\text{AR}}/I_{\text{CR}}$, which was then interpreted as a proxy for the ratio between the concentrations of RNA inside and outside the aggresome, $\phi_{\text{R,A}}/\phi_{\text{R,C}}$, with a prefactor $B$. Thus, to compare our model to the data, we fit

$$\frac{I_{\text{AR}}}{I_{\text{CR}}} = B \exp(-N_{\text{R}} \Delta H/kT), \tag{14}$$

to the data with $B$ and $\Delta H/kT$ as fitting parameters using matlab/octave's `lsqnonlin` function. The fitting results are discussed in the main text.

### V. SUPPLEMENTARY NOTE 5: DYNAMICS OF DIFFUSION AND DEGRADATION

We hypothesise that the aggresome provides an environment where the RNA is protected from the degradation mechanism that takes place in the cytosol. To explore this, we adapted the steady-state model above into a dynamic model. The dynamics by which the length distribution of RNA evolves in the aggresome is determined can be described by a diffusion-limited process. See Schematics in Fig. 3.

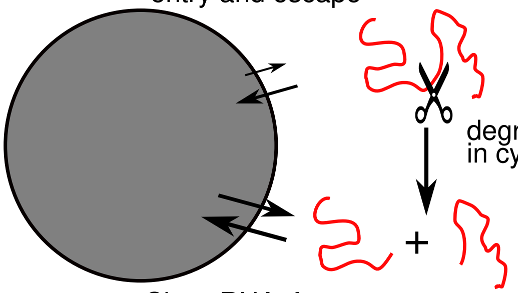


FIG. 3. Dynamics of RNA: the entry to and escape from the aggresome is limited by length-dependent diffusion, and degradation takes place in the cytosol. Part of figure accredited to Vecteezy.com.

Our model is similar to the Smoluchowski coagulation model [3], where the rate of diffusion-limited transport is controlled by the diffusivity and diffusive length scales inside/outside the droplets. However, while the coagulation model describes a change in droplet size, in our case the diffusion of RNA only alters the RNA concentration, but not the droplet size. Thus, we describe the rate by which a strand of length $N_{\text{R}}$ enters the aggresome using $K_{\text{diff,C}} = L_{\text{C}}^2/D_{\text{C}}$, with $L_{\text{C}}$ the characteristic diffusion length in the cytosol and $D_{\text{C}}$ the diffusivity. We assume the diffusivity to scale as $D_{\text{C}} \propto N_{\text{R}}^{\nu}$ based on mechanisms such as reptation ($\nu = -2$), Rouse diffusion ($\nu = -1$), and Zimm or Stokes-Einstein diffusion ($\nu = -1/2$). Here, we will assume the latter, and write $K_{\text{diff,C}} = K_{\text{diff}}^0 N_{\text{R}}^{-1/2}$. Similarly, the rate by which the RNA escapes the aggresome is set by the diffusivity and diffusion length within the aggresome, but modified by a Boltzmann factor that penalises the loss of associations of RNA with the contents of the aggresome, $K_{\text{diff,A}} = BK_{\text{diff}}^0 N_{\text{R}}^{-1/2} \exp(N_{\text{R}} \Delta H/k_{\text{B}}T)$. The rate coefficient $K_{\text{diff}}^0$ must be equal within and outside the aggresome in order to obey the detailed balance $K_{\text{diff,A}}/K_{\text{diff,C}} = B\phi_{\text{A}}/\phi_{\text{C}}$.

Thus, the dynamics by which RNA enters and escapes the aggresome can be described by the master equation

$$\frac{\partial}{\partial t}\phi_{\text{A}}(N_{\text{R}}, t) = K_{\text{diff}}^0 N_{\text{R}}^{-1/2} \phi_{\text{C}}(N_{\text{R}}, t) - BK_{\text{diff}}^0 N_{\text{R}}^{-1/2} \mathrm{e}^{-N_{\text{R}} \Delta H/k_{\text{B}}T} \phi_{\text{A}}(N_{\text{R}}, t), \tag{15}$$

with $\phi_\mathrm{A}(N_\mathrm{R}, t)$ the volume fraction of RNA with length $N_\mathrm{R}$ in the aggresome at time $t$, and $\phi_\mathrm{C}(N_\mathrm{R}, t)$ that in the cytosol.

The dynamics of the RNA in the cytosol is affected by degradation. While this can be treated in a sequence-dependent way, we will for simplicity use a mean-field approach where each bond between nucleotides may break with rate $K_\mathrm{deg}$. This gives

$$\frac{\partial}{\partial t}\phi_\mathrm{C}(N_\mathrm{R}, t) = -K^0_\mathrm{diff} N_\mathrm{R}^{-1/2}\phi_\mathrm{C}(N_\mathrm{R}, t) + BK^0_\mathrm{diff} N_\mathrm{R}^{-1/2} \mathrm{e}^{-N_\mathrm{R}\Delta H/k_\mathrm{B}T}\phi_\mathrm{A}(N_\mathrm{R}, t) \tag{16}$$

$$-(N_\mathrm{R}-1)K_\mathrm{deg}\phi_\mathrm{C}(N_\mathrm{R}, t) + 2\sum_{n=N_\mathrm{R}}^{\infty} K_\mathrm{deg}\phi_\mathrm{C}(n, t), \tag{17}$$

where the third term in the right-hand side of the equation describes the loss of long chains, and the fourth term describes the resulting generation of short chains.

As initial conditions we will assume all material resides in the cytosol at time $t = 0$, i.e., $\phi_\mathrm{A}(N_\mathrm{R}, t = 0) = 0$ and $\phi_\mathrm{C}(N_\mathrm{R}, t = 0) = \phi_\mathrm{tot}(N_\mathrm{R})$. To estimate the shape of the length distribution $\phi_\mathrm{tot}$ we use the screentape data in Fig 2a. From our fit of the intensity ratio in Eq. (14) to this data (displayed in Fig 2b), and by comparison to the concentration ratio in Eq. (11), we have estimated the total RNA concentration as $\phi_\mathrm{tot} \propto B\phi_\mathrm{A} + \phi_\mathrm{C}$. We describe this distribution empirically using

$$\phi_\mathrm{tot}(N_\mathrm{r}) \propto N_\mathrm{r} \exp\left(-10^{-6}(N_\mathrm{r} + 600)^2\right). \tag{18}$$

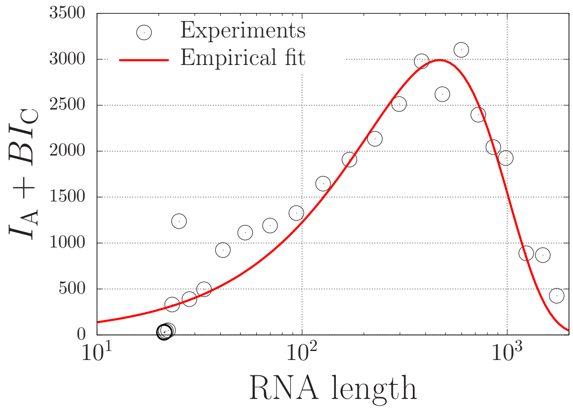


FIG. 4. Measured intensity of RNA in ($I_\mathrm{A}$) and outside ($I_\mathrm{C}$) the aggresome summed using weight $B$ to represent the total concentration $\phi_\mathrm{tot} \propto I_\mathrm{A} + BI_\mathrm{C}$ according to Eq. (14) and Eq. (11). The symbols are the ScreenTape data in Fig 2a (main text) using $B = 0.2457$ as determined by Fig 2b (main text) and the curve is the empirical fit given by Eq. (18).

The only free physical parameters in the dynamical equations are the diffusion and degradation rates, $K_\mathrm{diff}$ and $K_\mathrm{deg}$. By expressing time in units of the degradation rate, $K_\mathrm{diff}/K_\mathrm{deg}$ is the only physical parameter, which indicates that the system may be in crossover region between a diffusion- or degradation-limited regime that determine the time-dependence of the mean RNA length,

$$\langle N_\mathrm{R}(t)\rangle = \sum_{n=0}^{\infty} n(B\phi_\mathrm{A}(n, t) + \phi_\mathrm{B}(n, t)). \tag{19}$$

In our calculations we have used $B = 0.2457$ and $\Delta H = 0.0059 k_\mathrm{B}T$ as determined from the fit in Fig 2b.

We display representative time-dependent length distributions of the RNA in the cytosol and in the aggresome in the left panel of Fig. 5. At time zero, all material resides in the cytosol (top left panel). As time progresses, this peak decays because material diffuses towards the aggresome (top bottom panel). Subsequently, the material present in the cytosol degrades, and the distribution shifts to short RNA lengths in the cytosol, which may also diffuse to the aggresome, and may (depending on the ratio $K_\mathrm{diff}/K_\mathrm{def}$) lead to a bimodal distribution (bottom left panel).

The time-evolution of the mean RNA length, $\langle N_\mathrm{R}(t)\rangle$, is displayed in the right panel of Fig. 5 for a range of $K_\mathrm{diff}/K_\mathrm{deg}$ values. For slow diffusion rates, the RNA degrades before it can reach the aggresome. The typical timescale of RNA decay is length dependent, $(N_\mathrm{R}K_\mathrm{def})^{-1}$. That is, long RNA degrades relatively fast because of

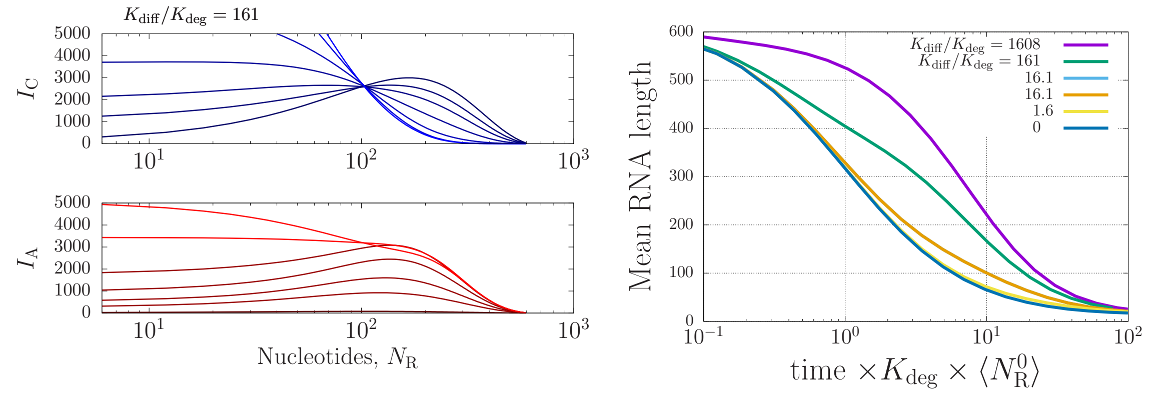


FIG. 5. Left: Calculated intensities within cytosol (blue; top) and within aggresome (red;bottom) for $K_{\rm diff}/K_{\rm deg} = 161$. The increasing brightness of the curves indicates more time has evolved. While long chains in the cytosol rapidly degrade (top panel), those that diffuse to the aggresome remain intact for long time scales until they escape the aggresome (bottom panel). Right: the mean chain length is measured as a function of time for various values of $K_{\rm diff}/K_{\rm deg}$. The time is given in units of $K_{\rm deg} \times \langle N_R^0 \rangle$, with $\langle N_R^0 \rangle$ the mean chain length at time $t = 0$.

our mean-field assumption, where long chains have a large number of bonds that can potentially break. However, independent of the sequence, the rate of degradation is delayed by the presence of the aggresome if diffusion is fast enough for RNA to reach the aggresome before it can degrade. This regime is diffusion-limited, because the time scale by which the RNA may degrade is determined by the time required to escape the aggresome. This time is set by the diffusion rate inside the aggresome, $K_{\rm diff} N^{-1/2}$, delayed by the length-dependent Boltzmann factor $\exp(N_R \Delta H / k_B T)$. This indicates that an increase of the association enthalpy $\Delta H$ delays the degradation rate; this may be achieved by aggresome compaction.

---

[1] Y. Shin and C. P. Brangwynne, Science **357**, eaaf4382 (2017), https://www.science.org/doi/pdf/10.1126/science.aaf4382.
[2] C. Schaefer, J. J. Michels, and P. van der Schoot, Macromolecules **49**, 6858 (2016), https://doi.org/10.1021/acs.macromol.6b00537.
[3] D. J. Aldous, Bernoulli **5**, 3 (1999).
